# Adversarial and variational autoencoders improve metagenomic binning

**DOI:** 10.1101/2023.02.27.527078

**Authors:** Pau Piera Líndez, Joachim Johansen, Arnor Ingi Sigurdsson, Jakob Nybo Nissen, Simon Rasmussen

## Abstract

Assembly of reads from metagenomic samples is a hard problem, often resulting in highly fragmented genome assemblies. Metagenomic binning allows us to reconstruct genomes by regrouping the sequences by their organism of origin, thus representing a crucial processing step when exploring the biological diversity of metagenomic samples. Here we present Adversarial Autoencoders for Metagenomics Binning (AAMB), an ensemble deep learning approach that integrates sequence co-abundances and tetranucleotide frequencies into a common denoised space that enables precise clustering of sequences into microbial genomes. When benchmarked, AAMB presented similar or better results compared with the state-of-the-art reference-free binner VAMB, reconstructing ∼7% more near-complete (NC) genomes across simulated and real data. In addition, genomes reconstructed using AAMB had higher completeness and greater taxonomic diversity compared with VAMB. Finally, we implemented a pipeline integrating VAMB and AAMB that enabled improved binning, recovering 20% and 29% more simulated and real NC genomes, respectively, compared to VAMB with moderate additional runtime. AAMB is freely available at https://github.com/RasmussenLab/VAMB.

## Introduction

It has been estimated that there are about one trillion (10^12^) species of microbes on Earth and that the large majority of these have yet to be discovered [1]. Decades ago, the standard approach to studying novel microbes was to isolate and cultivate them in the lab [2]. However, microorganisms establish and rely upon complex ecosystems that are not feasible to replicate in ex-natural environment conditions. This so-called “ cultivation bottleneck” limits culturing approaches to studying microbes [1]. In contrast, metagenomics enables culture-free microbial diversity characterization by analysing the entire set of genomes present in a given environmental sample [3]. Unfortunately, despite advances in sequencing throughput and bioinformatic tooling, reconstructing high-quality genomes from short read sequenced metagenomics samples remains challenging [4].

In particular, it is still not feasible to assemble reads from shotgun sequences to contigs that each cover whole original source genomes; instead, recovered genomes are often fragmented into many smaller contigs [5]. To mitigate this limitation of assemblers, contigs can be clustered into bins that represent their source genomes in a process called binning.

Hundreds of thousands of Metagenome Assembled Genomes (MAGs) which have previously been binned are publicly available, allowing detailed investigations of diverse microbial communities [6]–[8]. Despite the fact that several binners such as MetaBAT2, MaxBin 2.0, CONCOCT, and Canopy have been developed, binning performances are still far from optimal [9]–[12]. Recently we have developed a deep learning based method, called VAMB, that leverages variational autoencoders (VAEs) to obtain superior performance compared to previous reference-free binners [13]. VAEs are composed of an encoder that transforms the input features into a latent distribution, and a decoder that samples from the distribution and attempts to reconstruct the input from the sample. VAEs are widely used due to their ability to represent a complex feature space into a continuous latent distribution, and their inherent sampling process makes them suitable as generative models [14].

VAMB uses a VAE to integrate input contig abundances and tetranucleotide frequencies (TNF) to a common latent representation that can be clustered to yield bins. The regularisation of the latent space is done using Kullback-Leibler divergence with respect to a prior distribution, in VAMB’s case the Gaussian unit distribution [13]. However, another autoencoder framework is the adversarial autoencoder (AAE) where the regularisation of the latent space is achieved using another neural network, hence the name “ adversarial” [15]. Previous work applying AAEs to images showed that AAE models could generate latent representations with sharper and better-confined clusters compared to VAEs [15]. We, therefore, hypothesised that the application of an AAE for metagenomics binning could improve on clustering of near-complete genomes from the latent space. Furthermore, the original AAE implementation used an additional categorical latent space alongside the continuous one and showed that the model learned to cluster the input by assigning each cluster to a categorical class [15], [16]. We hoped that when applying AAEs to metagenomic sequences, the AAE would likewise learn to assign each genome into a single categorical class in the categorial latent space.

Here, we present Adversarial Autoencoders for Metagenomic Binning (AAMB), an extension of our original VAMB program. AAMB leverages AAEs to yield more accurate bins than VAMB’s VAE-based approach. We apply AAMB to both synthetic and real metagenomic benchmark datasets and show that more high-quality genomes are recovered using AAMB compared to using VAMB or other binners and that the extra genomes expand the taxonomic diversity of recovered genomes. We also present a method for automatically merging VAMB and AAMB, and show that the resulting ensemble method AVAMB is superior to both VAMB and AAMB while requiring nothing extra from the user other than a moderate increase in compute power.

## Results

### An adversarial autoencoder for metagenomics binning

Inspired by the original AAE implementation, the AAE in AAMB uses both a continuous (termed z) and a categorical (termed y) latent space (**Figure 1a**). Therefore, AAMB is able to extract bins both by clustering z like VAMB does [13], and by extracting the bin label directly from y. This resulted in two sets of bins, AAMB(z) and AAMB(y), respectively. When we investigated the structure of the AAMB z space when applied to the CAMI2 short-read human “ toy” datasets (see Methods), we found that distances between contigs from the same genome tended to be smaller than distances between contigs from different genomes for all benchmark datasets (**Supplementary Figure 1**), a prerequisite for clustering into genomes. AAMB’s z space was more compact than the VAMB latent space, which we believe was due partly to AAMB’s ability to also encode information in y. When clustering z, it yielded somewhat worse bins than VAMB, giving a total of 7% fewer NC genomes compared to VAMB across the CAMI2 and MetaHIT datasets (**Figure 1b**). The clusters of AAMB(y) were likewise inferior to VAMB, giving on average 39% fewer NC genomes. The relative performance of AAMB(z) vs AAMB(y) was dataset dependent; AAMB(*z*) outperformed AAMB(*y*) on the CAMI2 Airways, Gastrointestinal, Oral, Skin, and Urogenital datasets, reconstructing between 47-102% more NC genomes. On the contrary, AAMB(*y*), outperformed AAMB(*z*), on the MetaHIT dataset, by reconstructing 164% more NC genomes (**Supplementary Table 1**). Interestingly, the MetaHIT dataset has been difficult for all the binners we have tested, including MetaBAT2, MaxBin2.0 and Canopy. This suggested to us that the z and y spaces contained encodings of different subsets of the total information in the input data.

**Figure 1.**
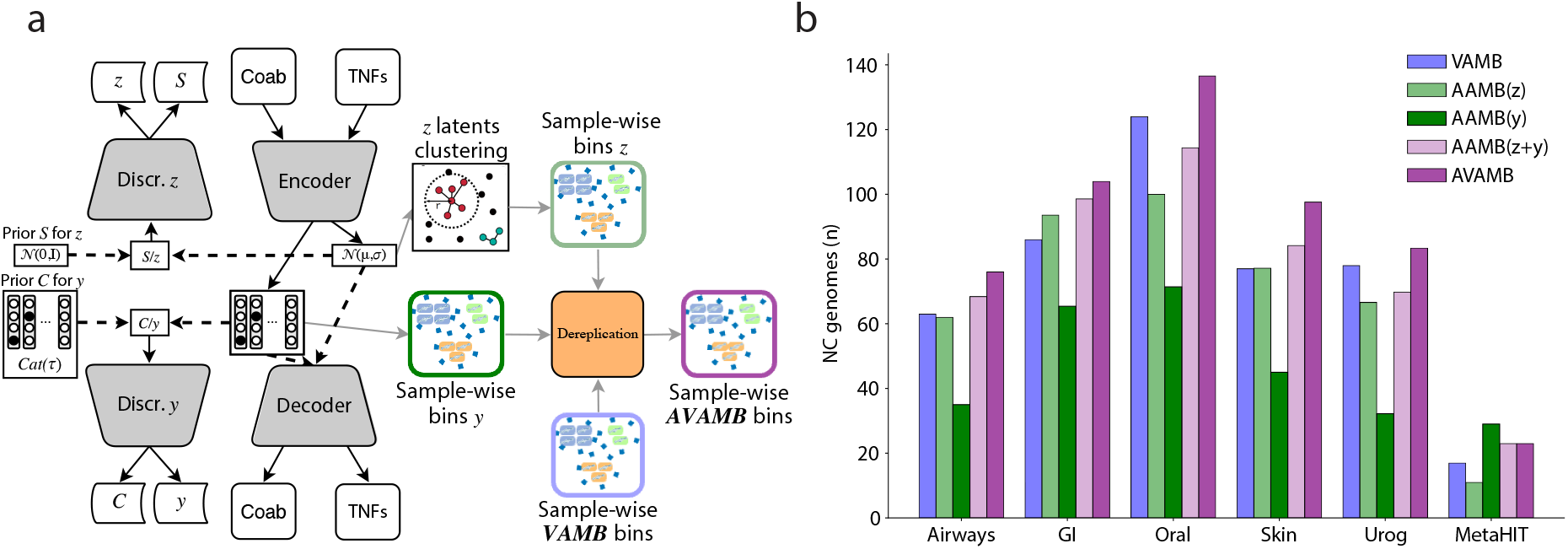
AAMB model overview and performance across the benchmark datasets. **A**. AAMB workflow overview. Tetranucleotide frequencies and abundances across samples are extracted per contig and input to the encoder. The encoder-decoder was optimised to reconstruct the input contig features from the regularized latent representations z and y. Regularisation is achieved by adversarial competition between the discriminators and the encoder, enforcing the latent encodings to stay close to their prior distributions. After training, latent representations z and y are retrieved. Then, the VAMB clustering algorithm was applied to generate clusters from the z latent representation, and cluster labels are taken directly from y. Finally, bins from z and y are deduplicated to the final AAMB clusters. These can then potentially be integrated with VAMB generated clusters. Dark arrows represent forward propagations. Dashed arrows represent sampling processes from the latent and priors, grey arrows represent clustering and dereplication steps performed after training AAMB. **B**. Number of distinct NC genomes reconstructed from the six benchmark datasets for VAMB (blue), AAMB(z) (light green), AAMB(y) (dark green), AAMB(z+y) (light purple), AVAMB (dark purple). GI, Gastrointestinal; Urog, Urogenital.

### The continuous and categorical latent space encodes different information

The original AAE paper showed how the y and z space primarily encoded high-level variance (“ class”) and low-level variance (“ style”), respectively. We hypothesised that AAMB behaved similarly, i.e. AAMB(y) would cluster contigs at a higher taxonomic rank than AAMB(z), which would imply that AAMB(y) might conflate lower taxonomic ranks. To test this, we measured the taxonomic distance between randomly selected contig pairs from the same cluster in AAMB(y) and AAMB(z) (**Supplementary Figure 2**). We found that AAMB(y) conflated higher ranks more often than AAMB(z), and AAMB(z) conflated strains more often than AAMB(y), which did not support our hypothesis. The hypothesis further implied that while most AAMB(z) clusters would have high strain-level purity, multiple AAMB(y) clusters representing different higher taxonomic ranks could sometimes be mapped to the same AAMB(z) space, causing some AAMB(z) clusters to be the union of otherwise pure strains from disparate high taxonomic ranks. If that was the case, it would imply that some contig pairs from the same AAMB(z) cluster would be from wildly different clades, but we did not observe this (**Supplementary Figure 3**). Further, when splitting AAMB(z) clusters by their y label to decontaminate this potential contamination, we found the resulting bins were no better (**Supplementary Table 2**), contrary to our hypothesis. We thus concluded that neither AAMB(z) nor AAMB(y) were redundant with respect to each other, nor did z only capture intra-y variance, but instead the two latent spaces learned complementary information. Therefore, we decided to explore the intersection and differences in genome reconstruction between VAMB, AAMB(z), and AAMB(y) (**Figure 2a**). We found that NC genomes reconstructed by AAMB(y) had a Jaccard index of 0.40 and 0.47 to the NC genomes reconstructed by AAMB(*z*), and VAMB, respectively (see Methods), while the Jaccard index of NC bins from AAMB(*z*) versus VAMB was higher at 0.64. Hence AAMB(z)’s genomes were more similar to VAMB’s than to AAMB(y)’s. This was not surprising, as the continuous Gaussian latent space of VAMB is more akin to the similarly structured z.

**Figure 2.**
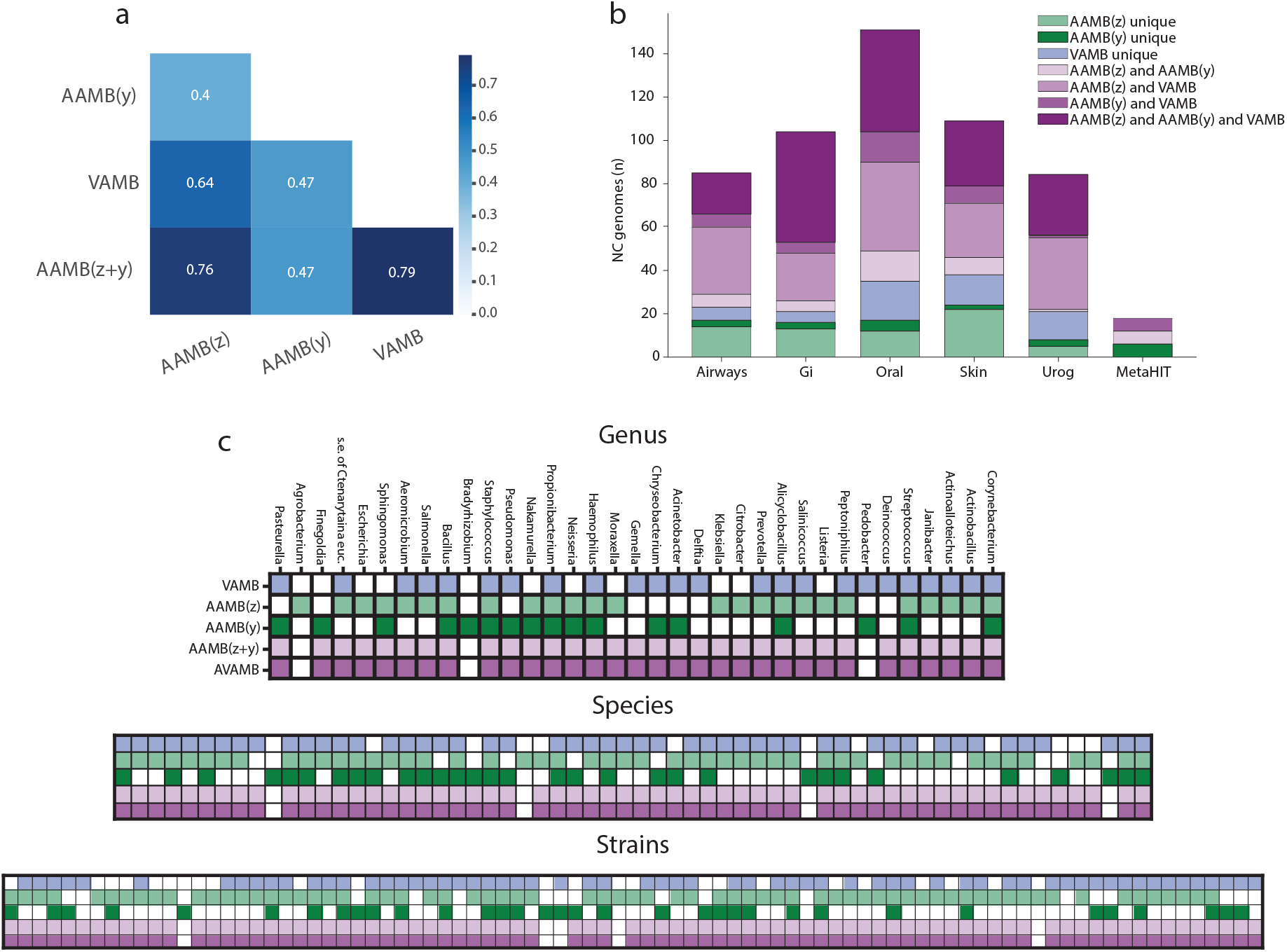
AAMB and VAMB reconstructed genomes analysis and integration with dRep. **A**. Jaccard correlation between the NC genomes from all benchmark datasets produced by AAMB(z), AAMB(y), VAMB, and AAMB(z+y) dereplicated bins. **B**. Contribution and the intersection of AAMB and VAMB for NC genomes on CAMI2 and MetaHIT datasets. The aggregated height of each bar expresses the total number of NC genomes reconstructed by VAMB and AAMB when dereplicating. unique: NC genomes only reconstructed by the given binner; and: NC genomes reconstructed by all the binner connected with the and operator and not by any other binner; NC: Near complete. **C**. Comparison of strains, species, and genera recovered at NC level by VAMB (blue), AAMB(z) (light green), AAMB(y) (dark green), as well as AAMB(z+y) (light purple), and AVAMB (dark purple). *s*.*e. of Ctenarytaina euc*., *secondary endosymbiont of Ctenarytaina eucalypti*.

### Combining the latent spaces AAMB(*y*) and AAMB(*z*)

Because AAMB(z) and AAMB(y) reconstructed different sets of genomes, we developed a technique to dereplicate the genomes sets reconstructed from the two latent spaces. Briefly, we assessed bin quality with CheckM2 [17] to remove low-quality bins, then for each bin pair that we deemed to be nearly identical, we removed the lowest-scoring bin. Finally, any contigs contained in two or more bins were assigned to the bin whose CheckM2 score would be improved the most by the addition of these contigs (see Methods). When comparing this dereplication method to a popular alternative MAG dereplication tool dRep [18], our method preserved more genomes, only losing 28 NC bins in the dereplication process across all the CAMI2 datasets, compared to 40 when using dRep (**Supplementary Table 3**). Simultaneously, our approach produced no duplicated contigs, whereas 6 contigs remain duplicated when using dRep (**Supplementary Table 4**). Applying this dereplication process to the union of AAMB(z) and AAMB(y), we obtained what we termed AAMB(z+y), and found that it outperformed VAMB, reconstructing 7% more NC genomes across all CAMI2 datasets (**Figure 1b**). This workflow came at some cost of computational power, as training and clustering time required were 1.9x and 3.4x higher when using GPU or CPU only, respectively, compared to VAMB (**Supplementary Table 5**). Because AAMB(z+y) had the best performance, we used it as the default AAMB workflow and will henceforth refer to it as simply AAMB.

### Combining the AAMB and VAMB framework

We found that AAMB (i.e. AAMB(z+y)) reconstructed a different set of NC genomes than VAMB (Jaccard index of 0.79) (**Figure 2b-c, Supplementary Figures 4-9**), and so hypothesised that we could apply the AAMB dereplication approach to also merge VAMB bins to form a VAMB+AAMB workflow we called AVAMB. The integrated AVAMB model proved highly performant as it allowed the reconstruction of 6-35% additional NC genomes compared to using only VAMB across the benchmark datasets (**Figure 1b, Supplementary Table 6**). Furthermore, few (2-7) NC genomes were lost during dereplication of AAMB and VAMB bins. We then investigated the contribution from AAMB(z), AAMB(y), and VAMB, and found that 23-50% of the NC genomes were reconstructed by all three methods across the benchmark datasets. On the other hand, we found that up to 27% of the NC genomes in a dataset were only identified by one method (**Figure 2b, Supplementary Table 7**). We concluded that VAMB, AAMB(z), and AAMB(y), each reconstruct different genomes from the benchmark datasets and that the dereplicated union of all three methods yields better overall performance. To check if the gains of AVAMB over VAMB were simply due to the former being an ensemble method, we merged the bins of VAMB and MetaBAT2 and compared this VAMB+MetaBAT2 with VAMB+AAMB. We found that VAMB+AAMB outperformed VAMB+MetaBAT2 on 5 of the 6 benchmark datasets, although an ensemble of all three methods did best on 5 of 6 datasets (**Supplementary Table 8**). This implies that AAMB has better synergy with VAMB than MetaBAT2. Since AAMB and VAMB share much of their pipelines including the computation of inputs to the encoders and are provided by the same software package, VAMB+AAMB is also significantly faster and easier to use than VAMB+MetaBAT2. Finally, we compared to using the semi-supervised binner SemiBin [19] and found that it reconstructed 15-36% more NC genomes compared to AVAMB on the CAMI data sets (**Supplementary Table 9**). This, however, also came at a higher computational cost taking 35-57 times longer than AVAMB (**Supplementary Table 10**).

### AAMB outperformed and complemented VAMB on real metagenomic datasets

The synthetic nature of the CAMI2 datasets enabled precise benchmarking, but because synthetic metagenomic datasets are not entirely realistic as they use idealised coverage distributions and read assemblies [4], we did not know if AAMB’s accuracy translated to real datasets. Therefore, we ran AAMB on two real datasets, a 1,000 gut microbiome dataset from Almeida et al. [20] and the Human Microbiome Project 2 Inflammatory Bowel Disease cohort with 1,306 longitudinal samples from 90 patients (HMP2 dataset) [21], and evaluated accuracy using CheckM2 (see Methods). We note that evaluating AAMB using CheckM2 was not entirely unbiased, since AAMB dereplicate similar bins using CheckM2 scores. Nonetheless, the observed results mirrored the ones seen on the synthetic CAMI datasets. AAMB(*z*) generated more NC bins than AAMB(y) with 1,464 and 1,963 more on the Almeida and HMP2 datasets, respectively. AAMB performed better than either AAMB(z) and AAMB(y) as well as better than VAMB, reconstructing 5,077 and 2,715 NC genomes, an increase of 9.7% and 2.3% (**Figure 3a-b**) compared to VAMB. Similarly, AVAMB did even better, yielding a total of 5,733 and 3,569 NC bins, which corresponded to an increase of 1,103 (24%) and 914 (34%) NC bins compared to VAMB. Similarly, we tried to use SemiBin on the Almeida data set, however, it did not complete within one week. From the samples that it finished (134), we estimated that it would take 52 days to complete using a GPU. In comparison, AVAMB finished in less than one day (22 hours).

**Figure 3.**
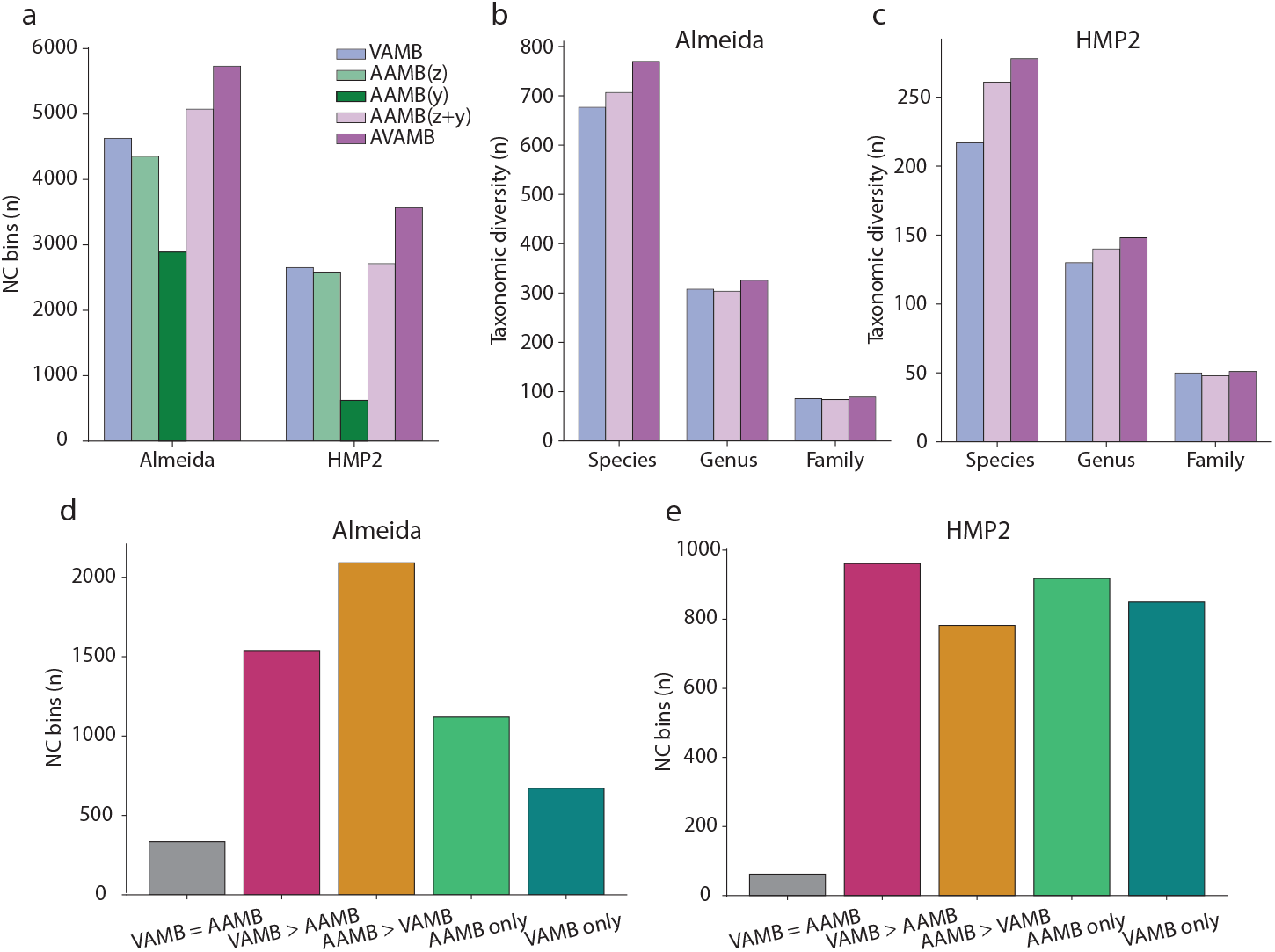
AAMB performance on real datasets. **A**. Number of NC bins reconstructed from the Almeida and HMP2 datasets for VAMB (blue), AAMB(*z*) (light green), AAMB(*y*) (dark green), AAMB(z+y) (light purple), AVAMB (dark purple). **B**. Number of taxa with at least one NC genome from the dereplicated set of bins generated by VAMB and AAMB on the Almeida dataset. AAMB and VAMB dereplicated bins were classified with GTDB-Tk. Taxa reconstructed only by VAMB (blue), taxa reconstructed only by AAMB (green), and taxa reconstructed by both AAMB and VAMB (purple). **C**. NC bins quality comparison between VAMB and AAMB. VAMB and AAMB bins were considered equal if belonging to the same sample with 100% identity over at least 75% of the smallest bin. NC: Near complete. V = A: VAMB and AAMB NC bins with exact same score. V > A: VAMB NC bins with a higher score than AAMB NC bins, A > V: AAMB NC bins with a higher score than VAMB NC bins, V unique: VAMB NC bins not reconstructed by any AAMB NC bin at the selected identity settings, A unique: AAMB NC bins not reconstructed by any VAMB NC bin at the selected identity settings.

### AVAMB recovers more distinct taxa than VAMB at higher quality

In order to investigate the nature of the additional genomes recovered by AAMB compared to VAMB, we assigned taxonomy to the NC bins recovered from the Almeida and HMP2 datasets (see Methods). We found that the additional genomes recovered by AAMB compared to VAMB increased diversity on the species level, but not higher taxonomic levels. As before, AVAMB performed better than either binner alone and increased diversity further on species level where it recovered 13% and 28% more unique species from the Almeida and HMP2 datasets respectively, compared to VAMB. Unlike AAMB, AVAMB also increased diversity on the genus level, where it reconstructed 5.5% and 14% more genera than VAMB (**Figure 3b**). To gain a better overview of which clades were improved using AVAMB compared to VAMB, we created a phylogenetic tree of core genes from NC bins recovered by AVAMB (**Figure 4**). In the figure, we marked bins that were uniquely recovered by AVAMB and not VAMB, and also marked bins that were recovered by both binners, but where the bin recovered by AVAMB was of better quality. AVAMB improved the quality of 2,091 NC bins also reconstructed by VAMB (**Supplementary Table 11**). We found that AVAMB recovered more bins across the entire tree with no apparent biases towards any particular clades, but that improved bins tended to cluster together in smaller clades across the tree.

**Figure 4.**
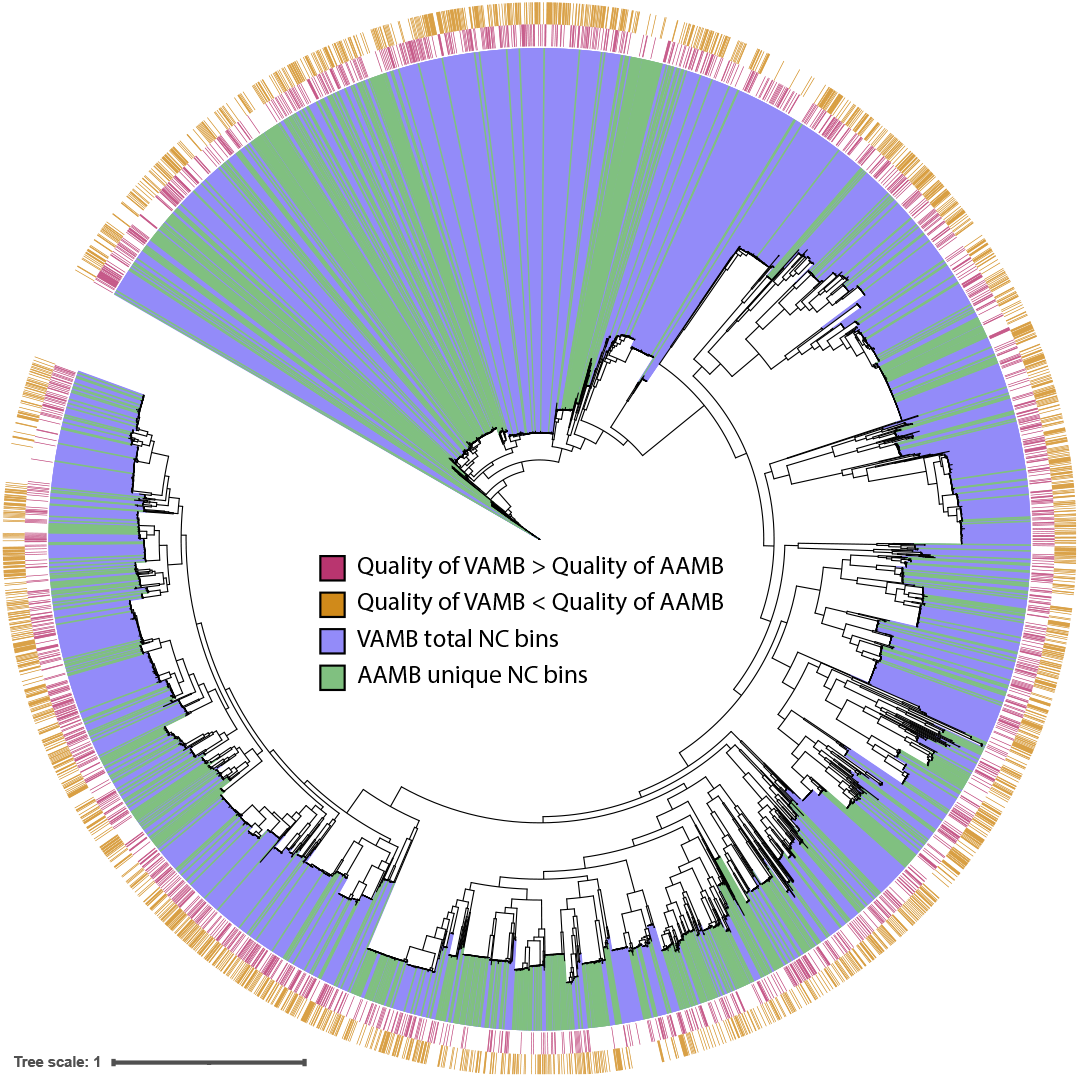
Maximum-likelihood phylogenetic tree of the NC bins reconstructed by VAMB and AAMB from the Almeida dataset. GTDB-tk was used to generate the multiple sequence alignment, IQ-tree to produce the tree, and iTOL to visualise the tree. Red stripe: bins reconstructed by VAMB with a higher score than AAMB, golden stripe: reconstructed by AAMB with a higher score than AAMB, blue range: bins reconstructed by VAMB, green range: bins reconstructed only by AAMB and not by VAMB.

## Discussion

We present AAMB, an adversarial autoencoder for metagenomic binning. AAMB is a probabilistic deep learning model that attempts to integrate and denoise contig features into two latent spaces. AAMB gave better results than VAMB, a state-of-the-art reference-free binning method. We found that AAMB bins excellently complemented VAMB bins in number, bin quality, and taxonomic diversity. Thus, dereplicating the union of VAMB and AAMB’s bins was found to maximise microbial genome recovery in both synthetic and real metagenomic datasets.

Dereplicating multiple binnings of the same contig set is not straightforward, but is simpler than dRep’s goal of dereplicating arbitrary genomes, because the former case involves duplication of the exact same contigs which is trivial to detect. Furthermore, pairs of imperfectly binned MAGs may have a nucleotide identity profile across the genome that is biologically unrealistic, with some large sections that are nearly identical, and other large sections with low nucleotide identity, a circumstance which dRep might not be built for. Hence, we believe the reason our dereplication method was more accurate than dRep was that our method was designed for the more narrow problem of dereplicating MAGs derived from the same set of contigs.

The increase in complexity of AAMB with respect to VAMB is not costless. We found that the training and clustering steps of the pipeline of AAMB were 1.8 to 2.7 times slower on average compared to VAMB’s when running on a GPU and CPUs, respectively. Furthermore, dereplication of AAMB and VAMB bins per sample must be accomplished if optimal bin integration is to be obtained, extending the total runtime by up to 4.2 minutes per sample for the largest datasets evaluated.

Like VAMB, AAMB leverages a multi-sample approach [13], and so is able to use its advantages, including more co-abundance signal from the same contigs, increased density of contig clusters by observing homologous contigs in multiple samples, and increased binning speed. Even while being slower than VAMB, AVAMB ran 28 times faster on the Almeida dataset than running MetaBAT2 on each sample and reconstructed 92% more NC bins. Expectedly, AVAMB reconstructed fewer NC genomes on the CAMI datasets than the semi-supervised binner SemiBin. However, this was also expected as being semi-supervised SemiBin uses database information to help bin the contigs at the cost of significantly increased runtime. On the contrary, AAMB and VAMB are unsupervised in their approach to generating the genome bins. Further improvement to AAMB, VAMB, and AVAMB could be done by implementing self or semi-supervised learning in the binning step itself in an efficient manner.

AAMB, the dereplication workflow, and AVAMB have been implemented using Python and Snakemake [22], and are shipped with the newest release of VAMB. From a software user perspective, it thus presents minimal differences with respect to the normal VAMB workflow.

## Methods

### Overview of AAMB

The AAMB workflow consists of the following main steps (**Figure 1a**). First, Tetra Nucleotide Frequencies (TNF) and per sample co-abundances (Coab) are extracted from the contigs and BAM files of reads mapped to contigs, and input to the AAMB model as a concatenated vector. The AAMB model encodes a continuous latent space (z) and a categorical latent space (y) and reconstructs the input from these two as the output. After training, two sets of contig clusters will be generated, one set from continuous latent space (z clusters) and another set from the categorical latent space (y clusters). The z space is clustered using the clustering algorithm of VAMB [13]. The y clusters are implicit from the one-hot categorical latent space and can simply be extracted as the categorical vector. Therefore, all contigs are present twice, both in the y and z output. The two sets were then split by sample of origin to per sample-specific bins using the principle of multi-split binning from the VAMB framework. Finally, the bins were filtered based on CheckM2 [17] scores and then dereplicated as described below.

### Datasets

We used the same synthetic benchmark datasets used in VAMB. For hyperparameter tuning, we used the short read Airways (n=10), Oral (n=10), and Urogenital (n=9) from the Critical Assessment of Metagenome Interpretation (CAMI2) short-read ‘toy’ human datasets [4]. Furthermore, we used the MetaHIT ‘error-free’ dataset as a training set as well [4]. The remaining two CAMI2 toy human datasets, CAMI2 Gastrointestinal (n=10), and CAMI2 Skin (n=10) were used for model validation. We further evaluated the methods on a 1,000-sample human gut microbiome dataset collected by Almeida et al. [20] and processed by Nissen et al. [25]. Additionally, we evaluated the methods on a dataset from the Human Microbiome Project 2 (HMP2) Inflammatory Bowel Disease (IBD) cohort consisting of 1,338 samples from a total of 27 healthy controls, 65 Crohn’s Disease, and 38 Ulcerative Colitis patients from Lloyd-Price et al. [21] and [21] processed by Johansen et al. [26].

### Contig pre-processing

Contig sequences were transcribed into numerical vectors following the same procedure described for VAMB [13] and summarised below. Both TNFs and co-abundances (Coab) were extracted for each contig. For the MetaHIT dataset [23], contig abundances were defined by the original authors whereas, for the CAMI2 [4], Almeida [20], and HMP2 [21] datasets the contig abundances were determined as done by Nissen et al [13]. In short, contigs for each sample were merged into a catalogue, and reads from each sample were aligned using minimap2 (v.2.15r905) [27] to the catalogue. For the HMP2 dataset, we used minimap2 version v.2.6. Abundances were calculated using the program *jgi_summarize_bam_contig_depths* from MetaBAT2 (v.2.10.2) [12, p. 2]. When input to VAMB and AAMB, the abundances were normalised across samples for the same contig to sum to one in order to mimic a probability distribution that a random mapping read will come from each sample. TNFs were calculated using the method as described in VAMB, i.e. projected into 103 orthonormal dimensions as originally described by Kislyuk et al. [28] and normalised by z-scaling each tetranucleotide across the contigs in order to increase the relative inter-contig variance [13]. As done in VAMB, each contig TNFs and Coab vectors were concatenated into a vector of number of samples + 103, constituting the model input vector [TNFs, Coab]^T^.

### Adversarial autoencoder architecture

The AAE architecture was composed of three modules: The encoder-decoder, the z latent space discriminator, and the y latent space discriminator, as done in the original work from Makhzani et al. [15] (**Figure 1a**). The encoder-decoder module learns the contig features and encodes them into the z and y latent spaces. The discriminators for the z and y latent spaces restrict the latents to be similar to their priors, thus performing the regularisation of the model, which imposes a structure on the latent space that makes it clusterable. The encoder is a sequence of two dense layers with 547 units dense layers, each with LeakyReLU activation function and batch normalisation. The encoder is connected to the dense layers μ, σ each with 283 units parameterizing z, and to the y layer with 700 units. Softmax activation is applied on the y latent layer to mimic a probability distribution. The decoder reconstructs the contig features using a sample of the Gaussian distribution z ∼ N(μ, σ), and the y vector. Its architecture is identical to the encoder. Considering that Coab was normalised across samples to sum to one, softmax was applied to the Coab output units. The two discriminators of the z and y space, D_z_ and D_y_, are both networks with the same architecture as the decoder model described above, except without batch normalisation, where the final output layer is one single node. Each discriminator is trained to discriminate between samples taken from the latent spaces (z, y) and the priors. The prior for the D_z_ is the unit Gaussian distribution N(μ=0, *σ*=I). For D_y_, the prior is the RelaxedOneHotCategorical [29] distribution Cat(*τ*), with the temperature *τ* = 0.15. The discriminators were optimised for a binary classification task, therefore the softmax activation function was used for the output node in order to interpret the output as a probability. All three modules are optimised with Adam [29], and are implemented using PyTorch (v.1.7.1) [29], with CUDA (v.8.0.61) being used when a GPU is available. For the results in this paper, we used an NVIDIA Tesla V100 GPU, and an Intel Xeon Gold 6230 CPU.

### Loss functions

The two discriminators were trained to minimise the difference between their binary prediction and the ground truth using binary cross entropy (BCE) on a pair of samples from the prior and the latent space, and hence use the loss:

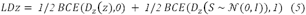

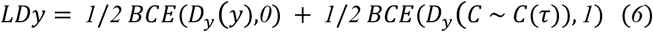

Where LDz and LDy is loss of the discriminators D_z_ and D_y_, respectively, and *S ∼ 𝒩*(*0, I*) and *C ∼ Cat*(*τ*) are samples from the priors, namely the standard normal distribution and the Gumbel-softmax distribution [30] with temperature parameter *τ*.

The encoder/decoder pair’s loss function is the sum of two terms: Reconstruction loss L_rec_ and regularisation loss L_reg_. L_rec_ encourages the networks to faithfully encode the input data’s information and was implemented as done in VAMB [13]. In short, the cross entropy (CE) loss was used for reconstructions of the abundances of the contigs (A_in_ vs A_out_) and mean squared error (MSE) for the reconstruction of the TNFs (T_in_ vs. T_out_). These two terms are weighted with hyperparameters w_coab_ and w_TNF_.

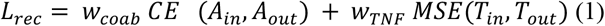

L_reg_ encourages the latent space to be similar to their priors and is the sum of two terms, which measure the ability of the discriminator to correctly identify the samples from the latent spaces:

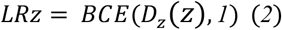

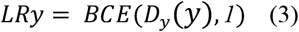

L_reg_ is then calculated as the sum of LRz and LRy, weighted by the slr hyperparameter:

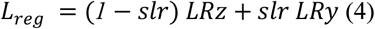

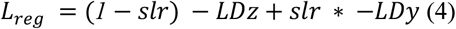

The final model loss L is computed by weighing L_rec_ and L_reg_ using the hyperpameter sl:

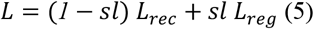

### Benchmarking

For the CAMI2 dataset, strains, species, and genus were defined exactly by the original authors [4]. For the MetaHIT dataset, strain definition was defined as the genomes used to generate the dataset, while species and genus were defined using NCBI taxonomy of these strains [24]. Calculation of AAMB bin quality within each taxonomy clade was done as in VAMB [13]: At the strain level, AAMB bin precision and recall were computed using the *vamb*.*benchmark* tools for all genomes in the dataset. Here *precision* quantifies the purity of the genome in a bin, and was defined as follows for pair (B, G) of bin B and genome G:

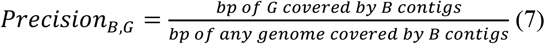

Similarly, recall quantify genome retrieval and was defined for bin B and genome G as follows:

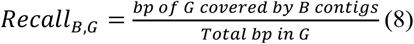

For a higher taxonomic clade L (e.g. species or genus), Recall_B,L_ and Recall_B,L_ was defined by

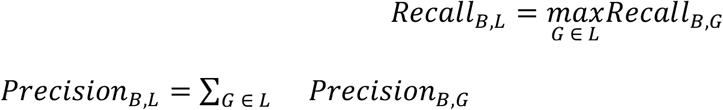

Binner performance is then given as the number of genomes reconstructed at some recall/precision threshold (typically 0.9/0.95) in any bin. SemiBin [19] v0.7.0 was run using the *multi_easy_bin* mode. In the run time comparison, we used 20 CPU, 1 GPU, 20 GB of RAM, for VAMB, AAMB, and AVAMB, and 1 CPU, 20 GPU, 150 GB of RAM for SemiBin.

### Hyperparameter searches

We did two random searches to select the hyperparameters of the AAMB model. During the first random search (RS1), the evaluated hyperparameters were selected to optimise the binning performance of the normally distributed latent space z (**Supplementary Figure 1**). Because adversarial models can be more unstable during training compared to variational models, the categorical latent y was not included. We first evaluated the reconstruction/regularisation scaling factor sl, and number/shape of the encoder/decoder hidden units, as these are important for stable competition between the encoder/decoder and the discriminators. Then, we added the categorial y latent, and optimised the related parameters slr and τ in a second random search (RS2), as these determine the model complexity for learning and encoding, respectively. (**Supplementary Figure 2**). For each iteration, the hyperparameters were randomly sampled within a given range, training the model independently on each of the training datasets: CAMI2 Airways, CAMI2 Urog, CAMI2 Oral and MetaHIT. Performance was evaluated based on the number of reconstructed genomes with precision above 0.9 and recall above 0.9. The final hyperparameters were sl = 0.0964, slr = 0.5, τ = 0.1596, with encoder and decoders of two hidden layers with 547 hidden nodes per layer, 283 nodes in the latent z layer, and 700 nodes for the categorical y layer. However, the categorical y layer dimension could be always adjusted depending upon the estimated taxonomic diversity. Thus, we decided to increment the y layer size for the Almeida and HMP2 datasets since Nissen et al. [13] reported a higher diversity with respect to the CAMI2 and MetaHIT datasets.

### De-replicating genomes between latent spaces

Because the latent spaces each encode every contig, the contigs are binned multiple times, and the same bin may be output multiple times in AAMB. Therefore, we devised a technique to dereplicate the sets of genomes. First, we ran CheckM2 [17] v0.1.3 to assign completeness and contamination to all genomes and removed all genomes that were not NC (i.e. completeness > 0.9, contamination < 0.05). Each genome was assigned a score computed as

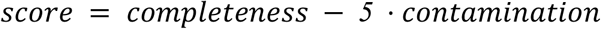

We then identified all “ near-identical” pairs of bins, where at least 75% of the smaller bin’s nucleotide content was present in the larger bin. For each of these pairs, we removed the bin with the lowest score. For each contig still shared by multiple bins, we created a bin without that contig and scored it with CheckM2 as above. The contig was then assigned the bin to which its removal would result in the largest score drop. To compare this technique against dRep [18], we ran dRep v3.0.0 on each sample independently using default parameters.

### AAMB and VAMB complementarity tree

NC bins from AVAMB were dereplicated and scored as described above. If a pair of bins, one from VAMB and the other from AAMB was identified as near-identical as above, then we consider the retained of the two bins to be recovered from both AAMB and VAMB. From the dereplicated AVAMB bins, we ran GTDB-tk [31] v.2.1.0 on the bins both to assign taxonomies to each bin, and to obtain a multiple sequence alignment, from which we inferred a phylogenetic tree under the LG amino acid substitution model using IQ-TREE [32] v1.6.8. We annotated and visualised the tree with iTOL [33].

## Acknowledgments

P.P., J.J., J.N.N., A.I.S. and S.R. were supported by the Novo Nordisk Foundation grant NNF14CC0001. Furthermore, P.P., J.N.N., and S.R. were supported by the Novo Nordisk Foundation grant NNF20OC0062223. In addition, S.R. was supported by the Novo Nordisk Foundation grant NNF19SA0059348. Finally, this work was also supported by the Novo Nordisk Foundation grant NNF21SA0072102.

## Author contributions

S.R. conceived the study and guided the analysis. P.P. developed AAMB and AVAMB and wrote the software and performed the analysis. Additionally, J.N.N. also performed analyses and wrote the software. J.J. and A.I.S. provided guidance and input for the analysis. P.P., J.N.N., and S.R. wrote the manuscript with contributions from all co-authors. All authors read and approved the final version of the manuscript.

## Data availability

The sequence data used in this study are publicly available from the respective studies or ENA. The semisynthetic MetaHIT dataset was downloaded from https://portal.nersc.gov/dna/RD/Metagenome_RD/MetaBAT/Files/ as the files depth.txt.gz and assembly-filtered.fa.gz. The simulated CAMI2 datasets were downloaded from https://data.cami-challenge.org/participate from ‘2nd CAMI Toy Human Microbiome Project Dataset’. The Almeida *de novo* assemblies were downloaded from http://ftp.ebi.ac.uk/pub/databases/metagenomics/umgs_analyses/benchmarked_assemblies.tar.gz and the reads were downloaded from ENA as specified in their publication. The HMP2 data was originally obtained from the European Nucleotide Archive accession PRJNA398089 and assembled as described in Johansen et al., 2022.

## Code availability

All code for AAMB and VAMB, as well as the dRep workflow, can be found on GitHub at https://github.com/RasmussenLab/vamb and are freely available under the permissive MIT license.

