## Supplementary material for "Adversarial and variational autoencoders improve metagenomic binning"

**Author List Footnotes**

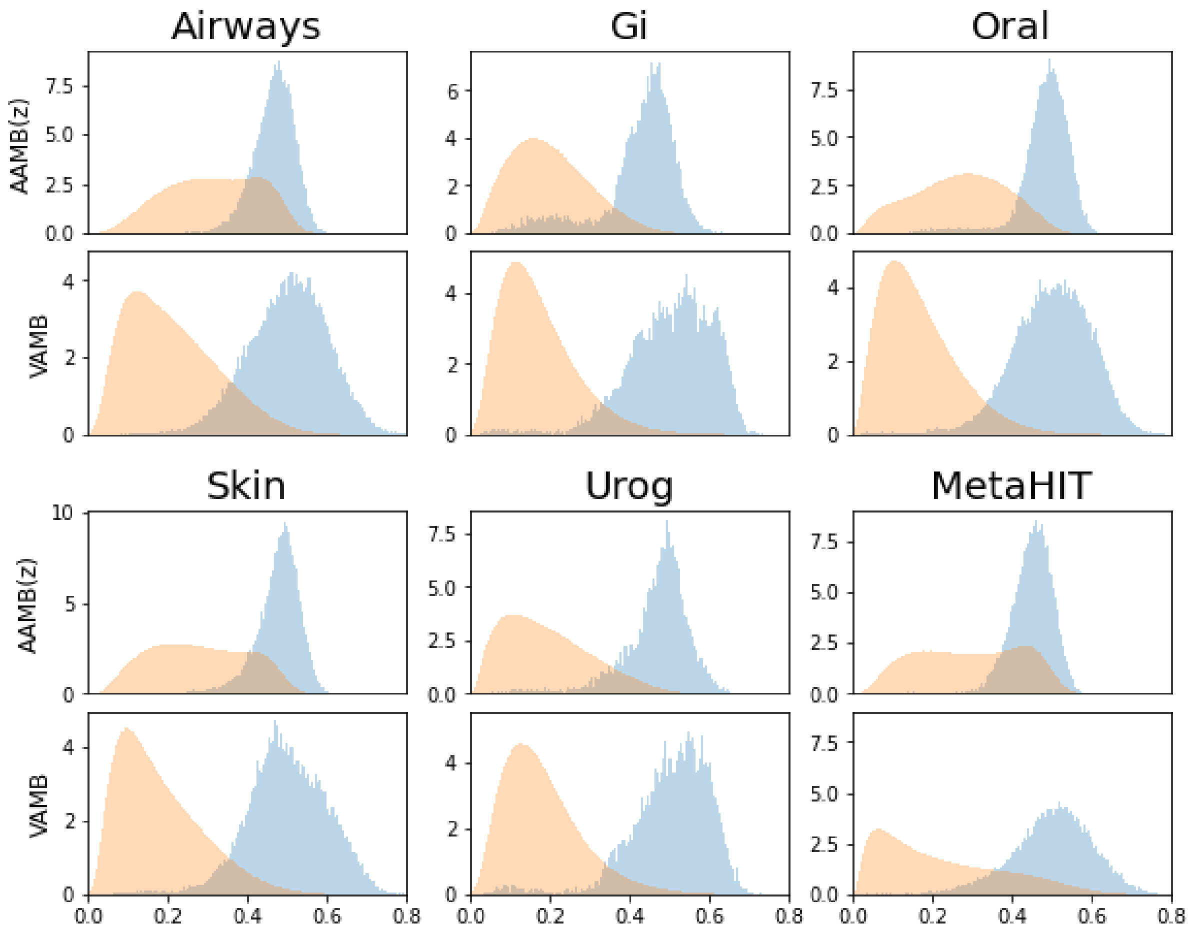


**Supplementary Figure 1. AAMB(z) and VAMB Intra-inter genome contig distance distributions when trained on CAMI2 and MetaHIT datasets.** Distribution of the distances of contigs from the same genome, i.e. intra genome contig distances, are plotted in orange. Distribution of the distances of contigs from different genomes, i.e. inter genome contig distances, are plotted in blue. Distributions are plotted for all CAMI2 and MetaHIT datasets, and for both AAMB(z) and VAMB. Airways, CAMI2 Airways; Gi, CAMI2 Gastro intestinal; Oral, CAMI2 Oral; Skin, CAMI2 Skin; Urog, CAMI2 Urogenital.

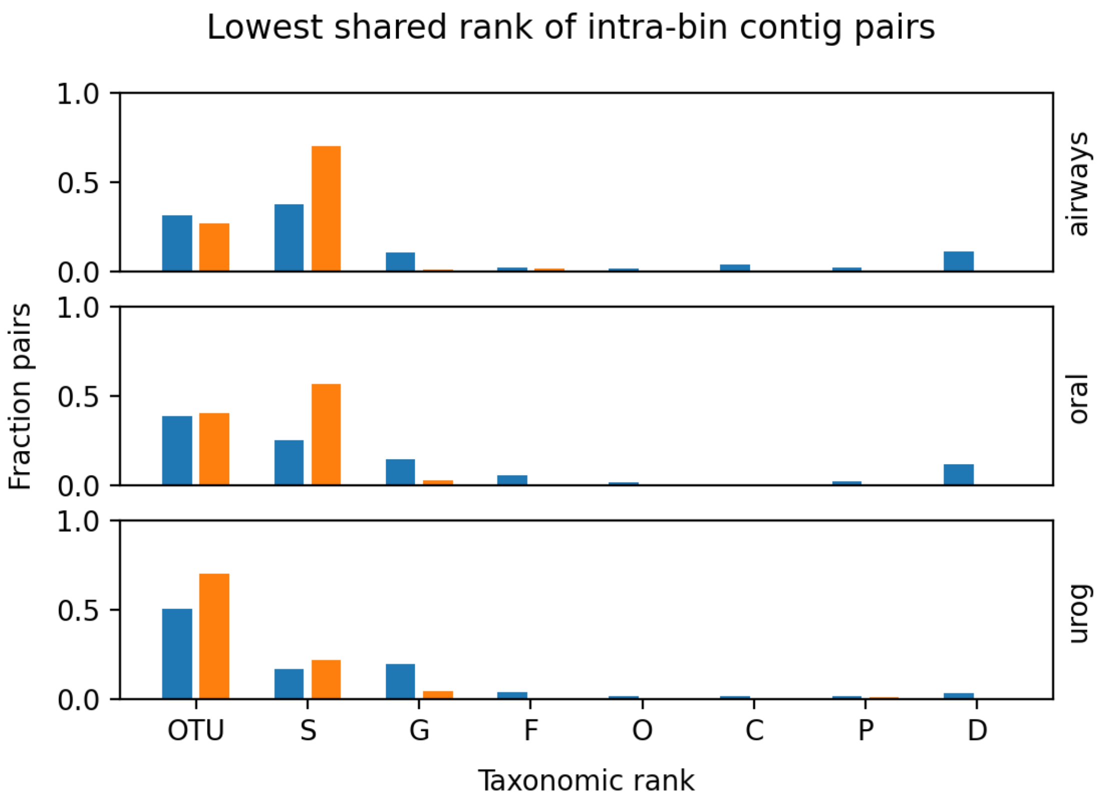


**Supplementary Figure 2. Phylogenetic distance between contigs from the same AAMB(y) or AAMB(z) cluster.** From each cluster in AAMB(y) (blue color) and AAMB(z) (orange color), we sampled 25 contig pairs, and computed the lowest shared taxonomic rank of each pair. The figure shows the distribution among sampled pairs.

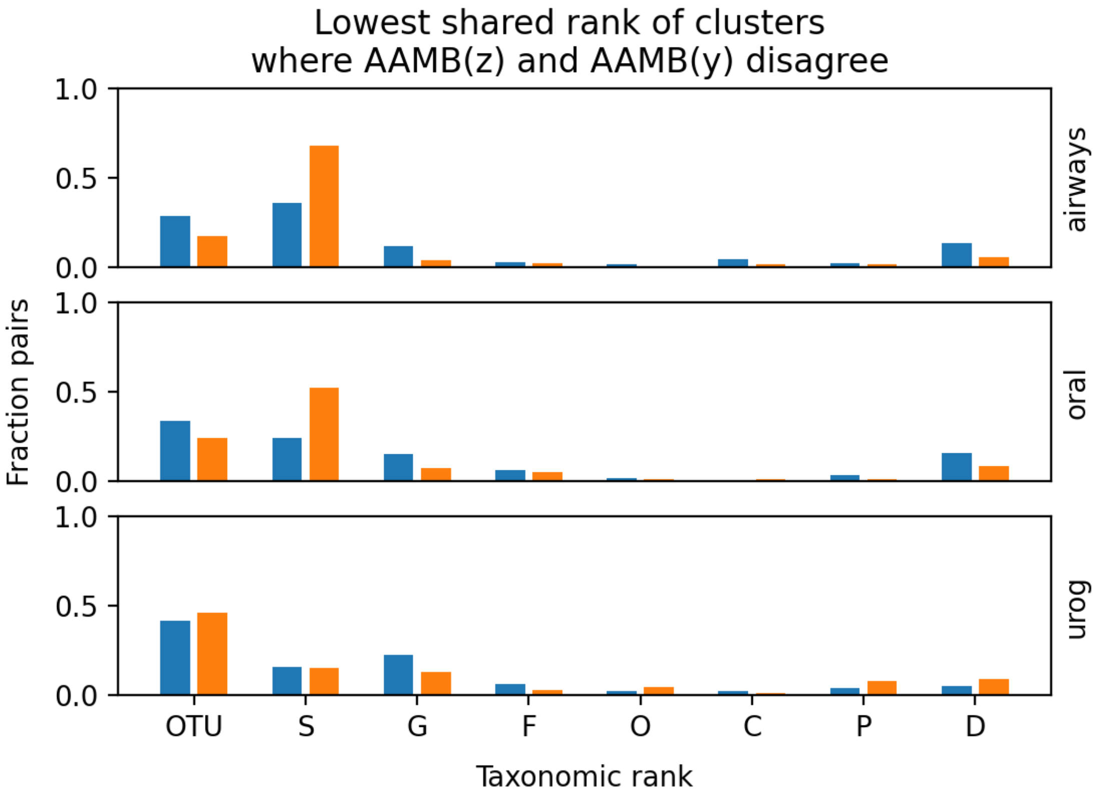

**Supplementary Figure 3. Phylogenetic distance between contigs pairs where AAMB(z) and AAMB(y) disagree on whether they belong to the same cluster.** Same as Figure 15, but here, within an AAMB cluster, we only sampled pairs of contigs that were placed in different AAMB clusters of the other latent space. Blue bars: Same AAMB(y), different AAMB(z). Orange bars: Same AAMB(z), different AAMB(y).

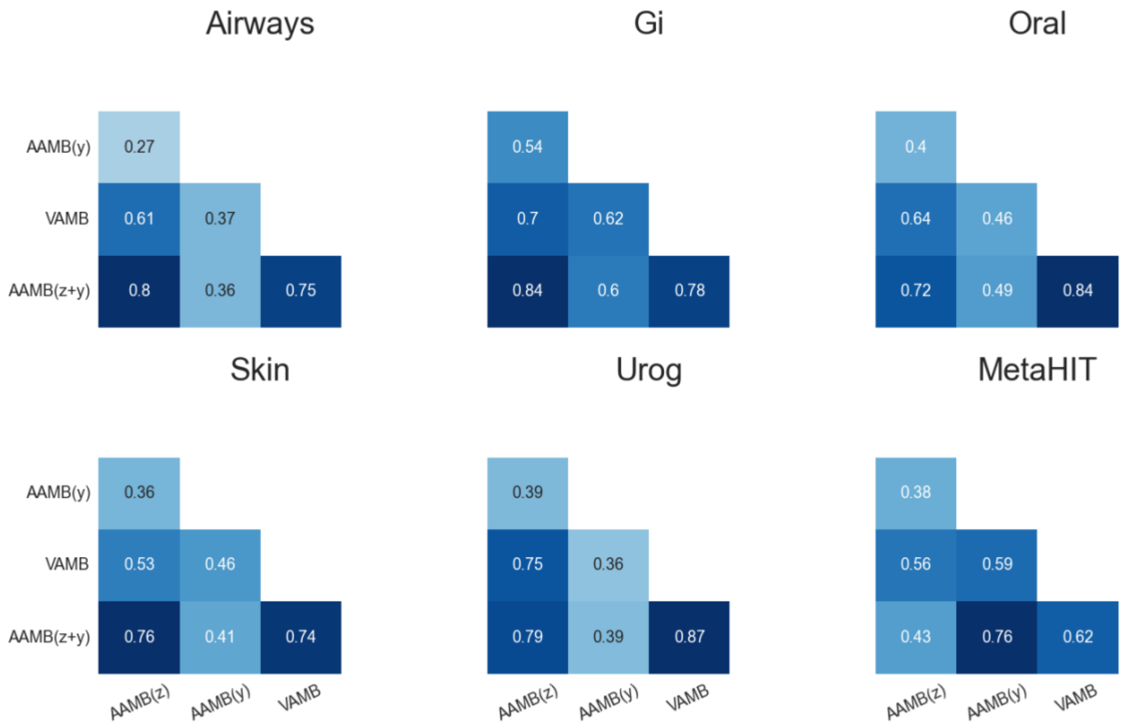


**Supplementary Figure 4. Jaccard correlation indexes between AAMB z, AAMB y and VAMB NC genomes along all benchmark dataset.** Jaccard correlation indexes between the NC genomes produced by AAMB *z* latent space, AAMB *y* latent space, VAMB, AAMB z latent space plus AAMB y latent space dereplicated bins, along all benchmark datasets. AAMB(z+y): dereplicated bins from AAMB z and AAMB y latent spaces; Airways, CAMI2 Airways; Gi, CAMI2 Gastrointestinal; Oral, CAMI2 Oral; Skin, CAMI2 Skin; Urog, CAMI2 Urogenital.

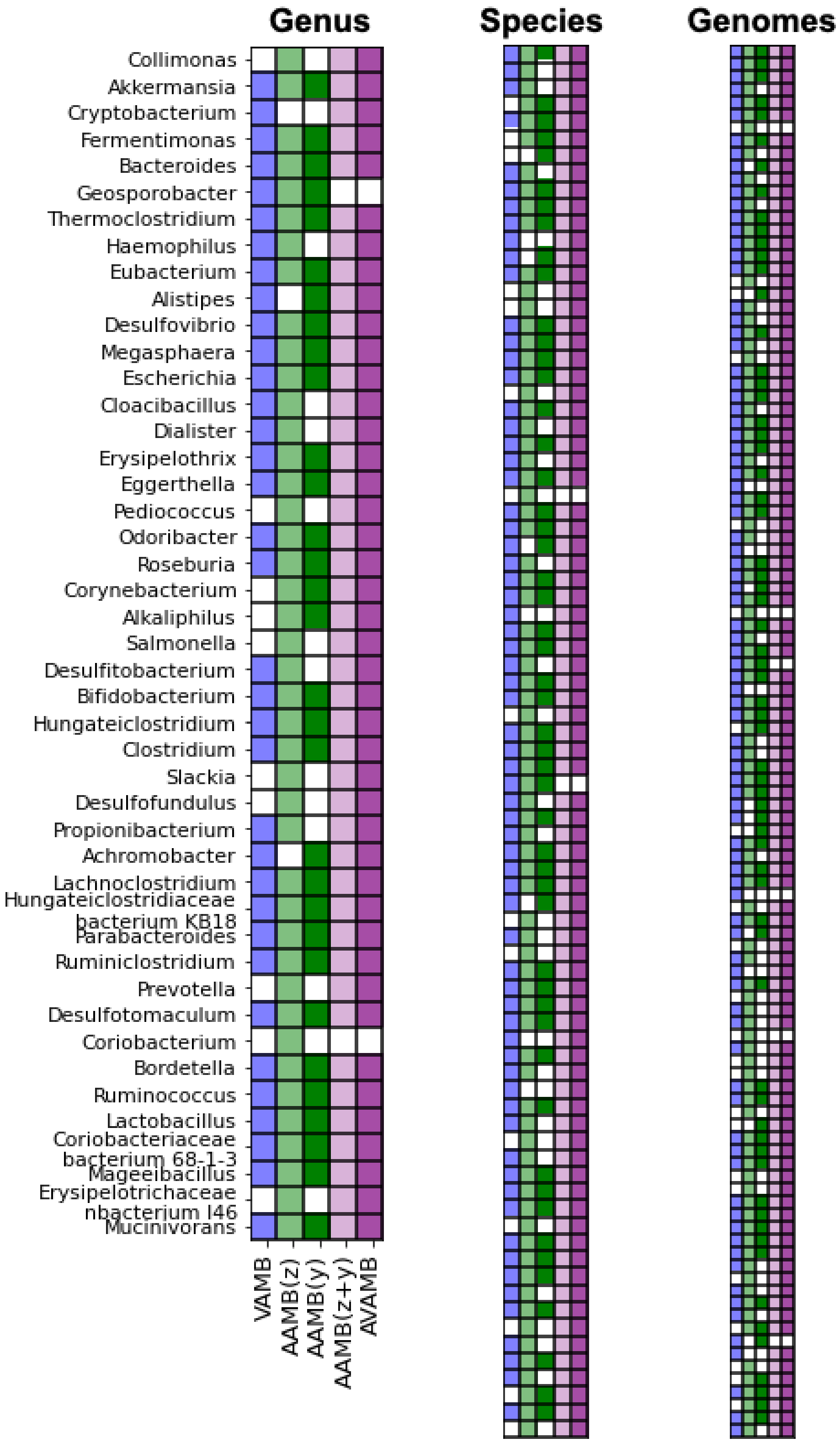


**Supplementary Figure 5. AAMB VAMB reconstructed genomes analysis and integration for the CAMI2 Gastrointestinal dataset.** Comparison of NC strains, species and genus sets of bins generated by VAMB (blue), AAMB(z) (light green), AAMB(y) (dark green), as well as the integrated dereplicated AAMB(z) plus AAMB(y) (light purple), AAMB(z) plus AAMB(y) plus VAMB dereplicated bins (dark purple). Gi, CAMI2 Gastrointestinal.

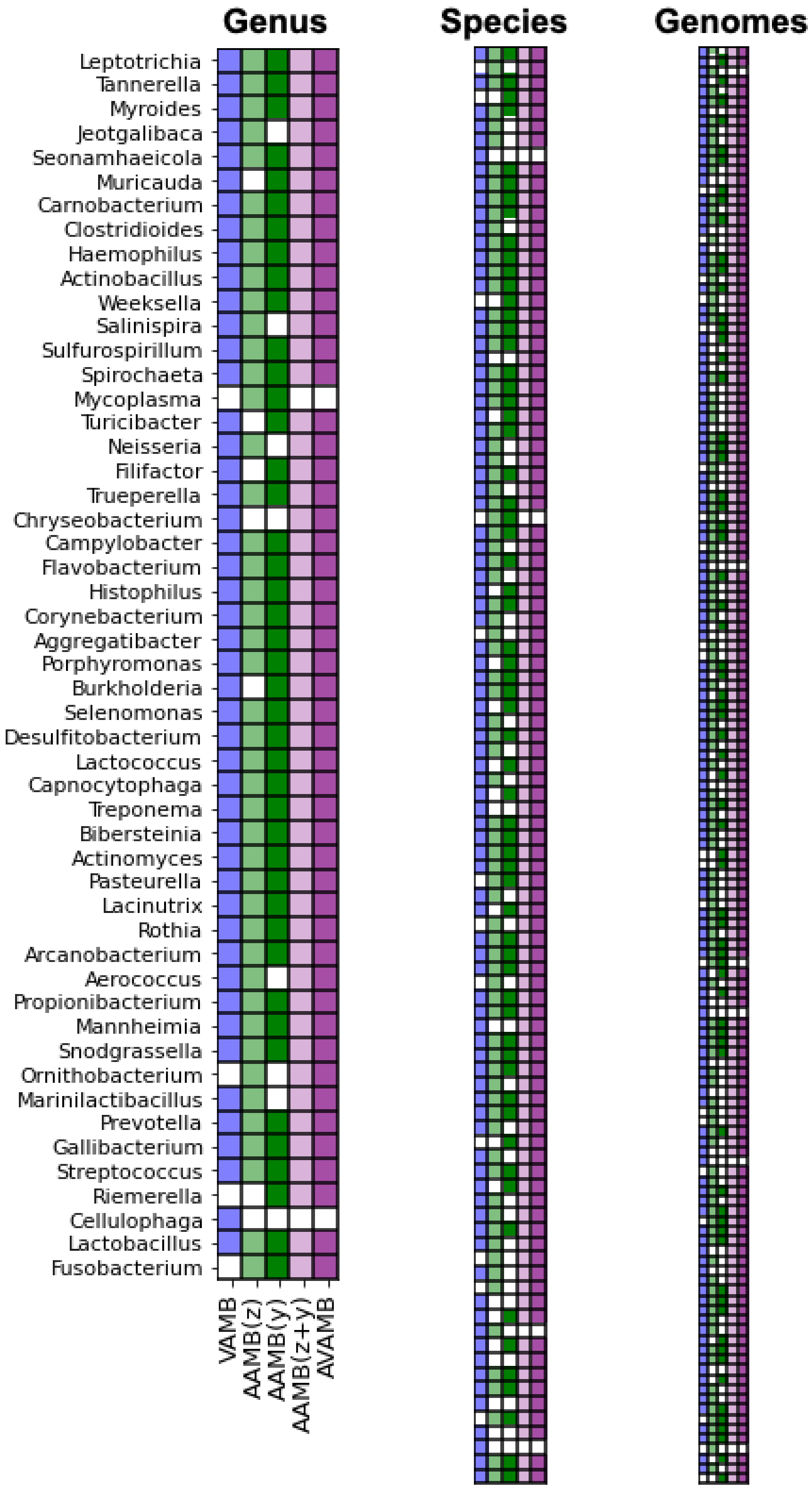


**Supplementary Figure 6. AAMB VAMB reconstructed genomes analysis and integration for the CAMI2 Oral dataset.** Comparison of NC strains, species and genus sets of bins generated by VAMB (blue), AAMB(z) (light green), AAMB(y) (dark green), as well as the integrated dereplicated AAMB(z) plus AAMB(y) (light purple), AAMB(z) plus AAMB(*y*) plus VAMB dereplicated bins (dark purple). Oral, CAMI2 Oral.

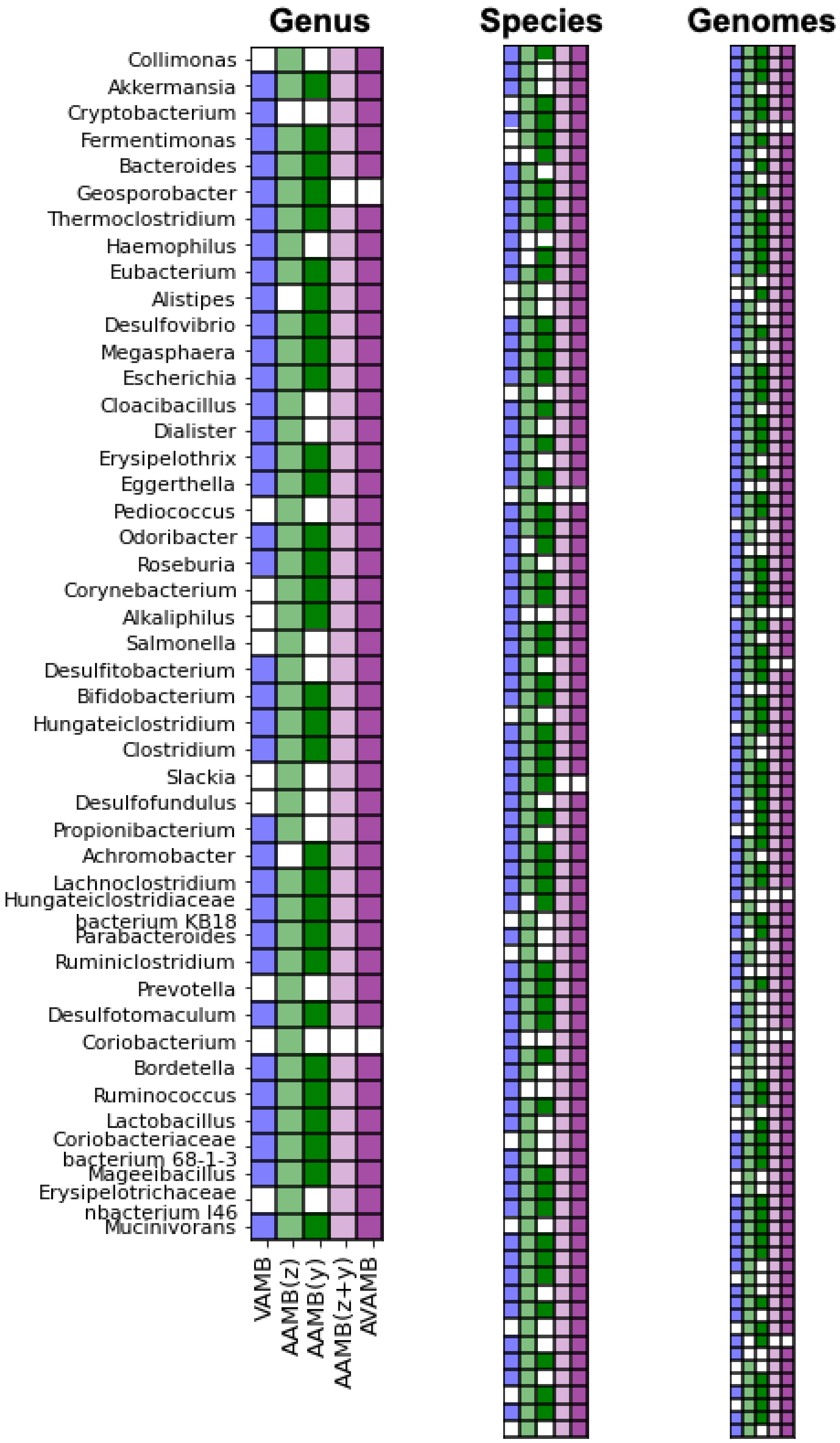


**Supplementary Figure 7. AAMB VAMB reconstructed genomes analysis and integration for the CAMI2 Skin dataset.** Comparison of NC strains, species and genus sets of bins generated by VAMB (blue), AAMB(z) (light green), AAMB(y) (dark green), as well as the integrated dereplicated AAMB(z) plius AAMB(y) (light purple), AAMB(z) plus AAMB(y) plus VAMB dereplicated bins (dark purple). Skin, CAMI2 Skin.

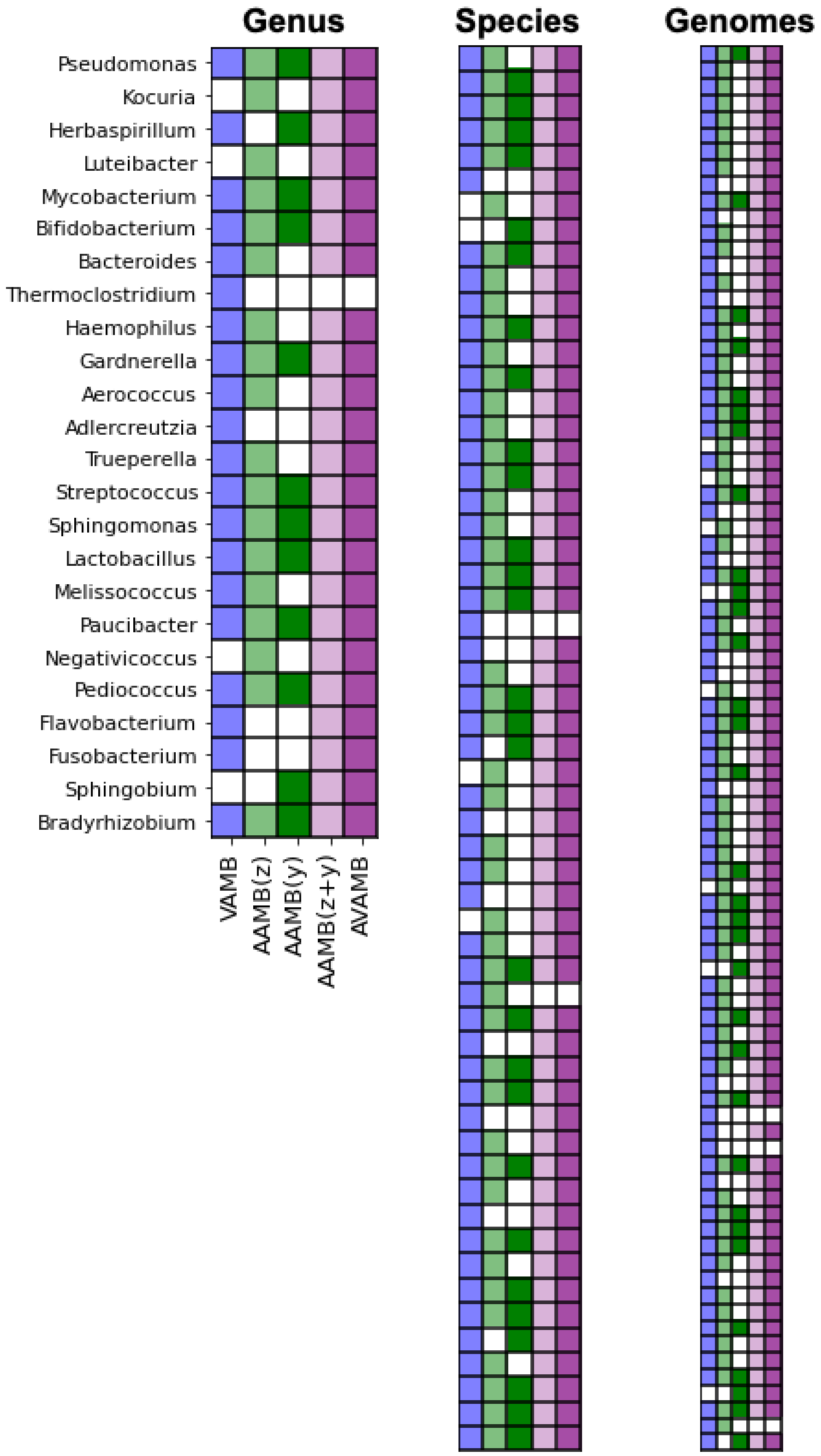


**Supplementary Figure 8. AAMB VAMB reconstructed genomes analysis and integration for the CAMI2 Urogenital dataset.** Comparison of NC strains, species and genus sets of bins generated by VAMB (blue), AAMB(z) (light green), AAMB(y) (dark green), as well as the integrated dereplicated AAMB(z) plius AAMB(y) (light purple), AAMB(z) plus AAMB(y) plus VAMB dereplicated bins (dark purple). Urogenital, CAMI2 Urogenital.

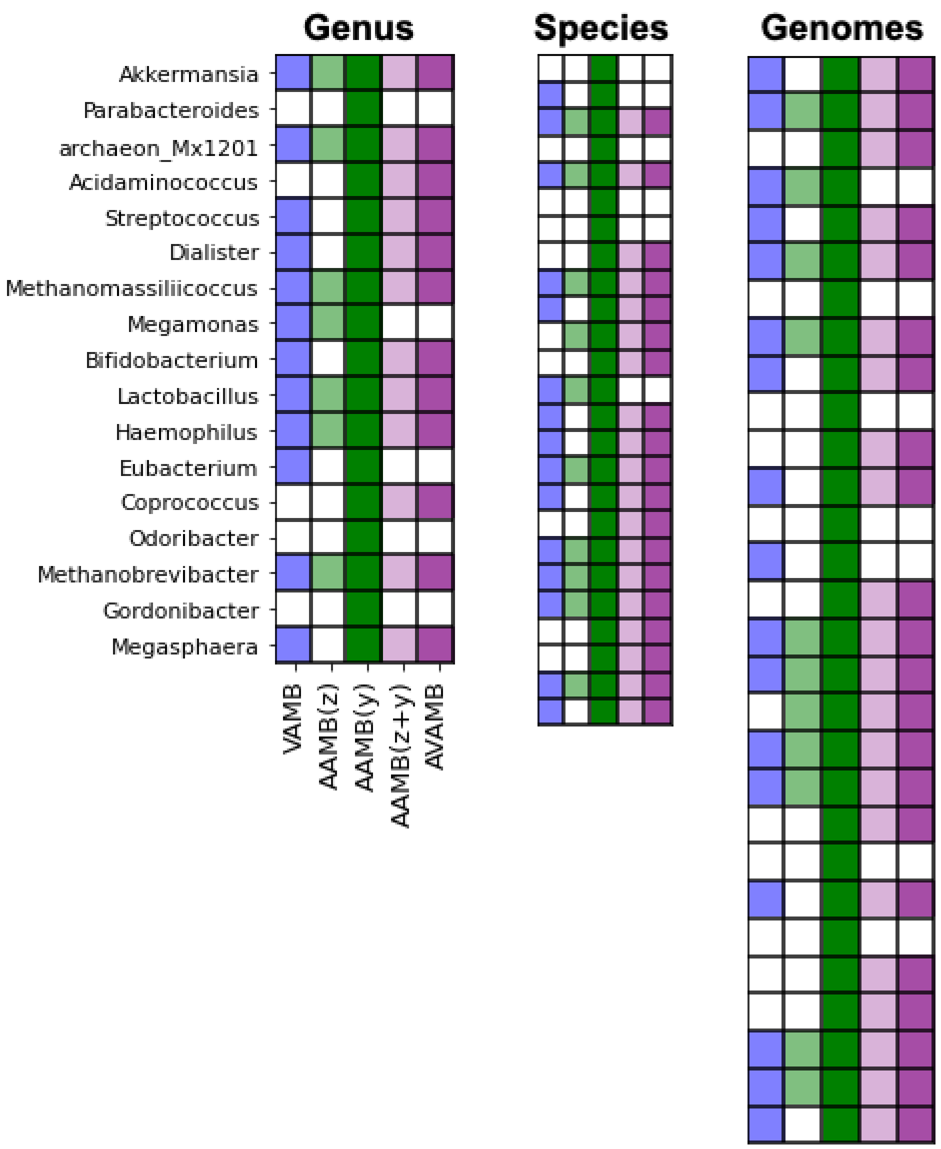


**Supplementary Figure 9. AAMB VAMB reconstructed genomes analysis and integration for the MetaHIT dataset.** Comparison of NC strains, species and genus sets of bins generated by VAMB (blue), AAMB(z) (light green), AAMB(y) (dark green), as well as the integrated dereplicated AAMB(z) plus AAMB(y) (light purple), AAMB(z) plus AAMB(y) plus VAMB dereplicated bins (dark purple).

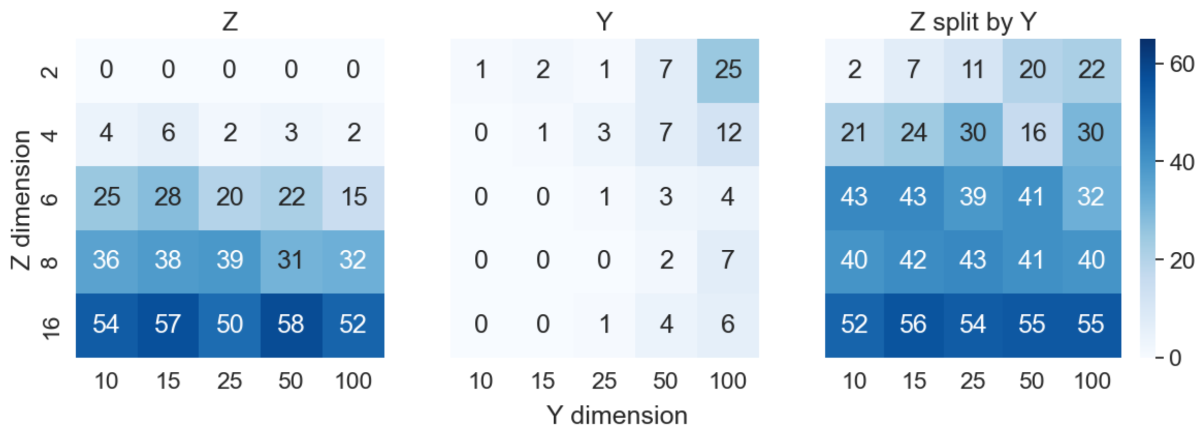


**Supplementary Figure 10. NC bins reconstructed from AAMB(z), AAMB(y) and AAMB(z) split by their AAMB(y) along z and y hyperparameter values.** For each z, y hyperparameters values, AAMB was run over the Airways CAMI2 dataset. For each run, NC bins were evaluated from the z latent space, y latent space and z latent space after splitting by y.

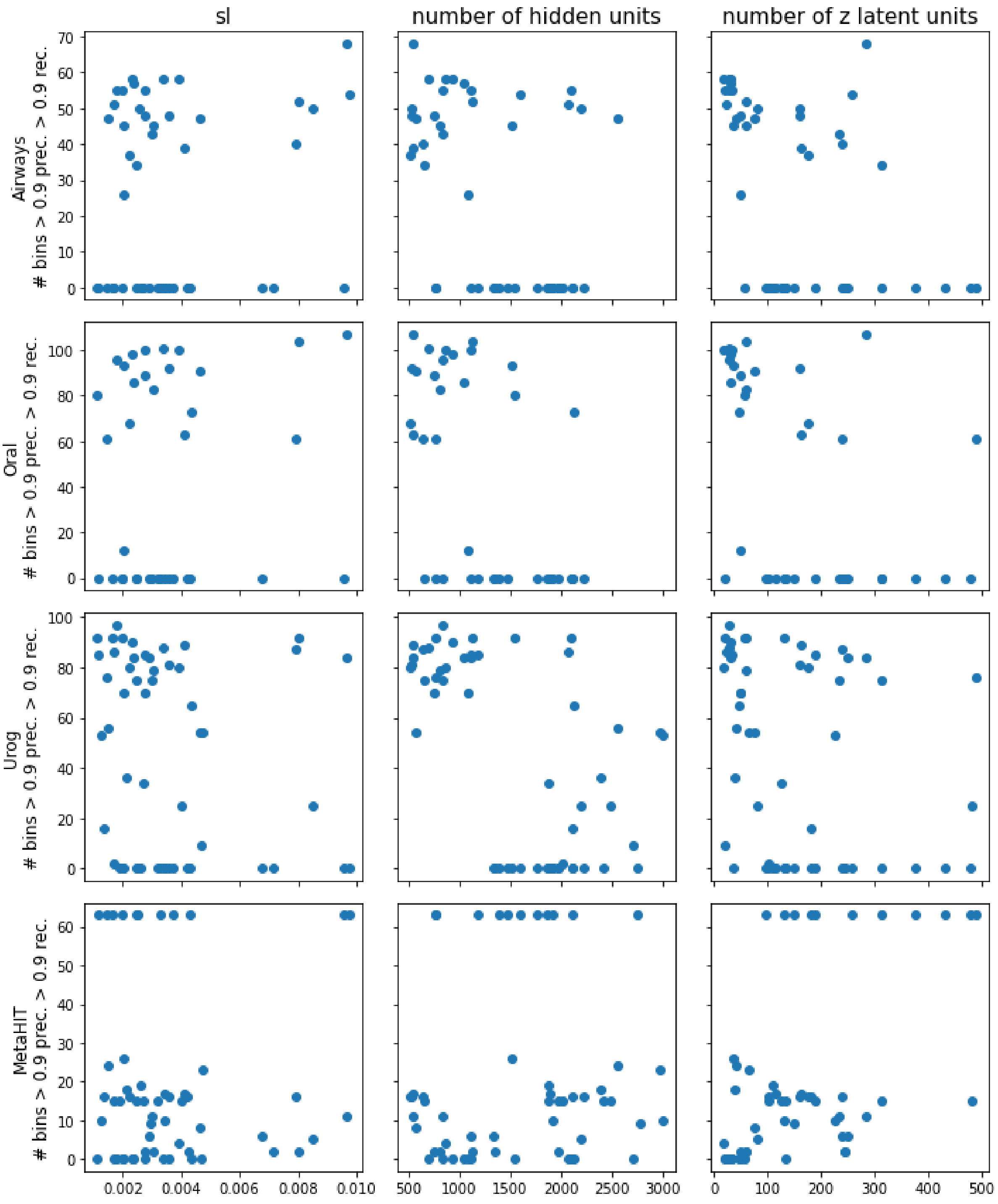


**Supplementary Figure 11. First round of hyperparameter random searches.** AAMB(z) bins above 0.9 precision and 0.9 recall are shown for Airways, Oral and Urogenital (Urog) CAMI2 datasets as well as for the MetaHIT dataset. Hyperparmeters evaluated are the loss scale (sl), the number of hidden units of the encoder-decoder (number of hidden units) and the number of latent units of the *z* latent space (number of latent units).

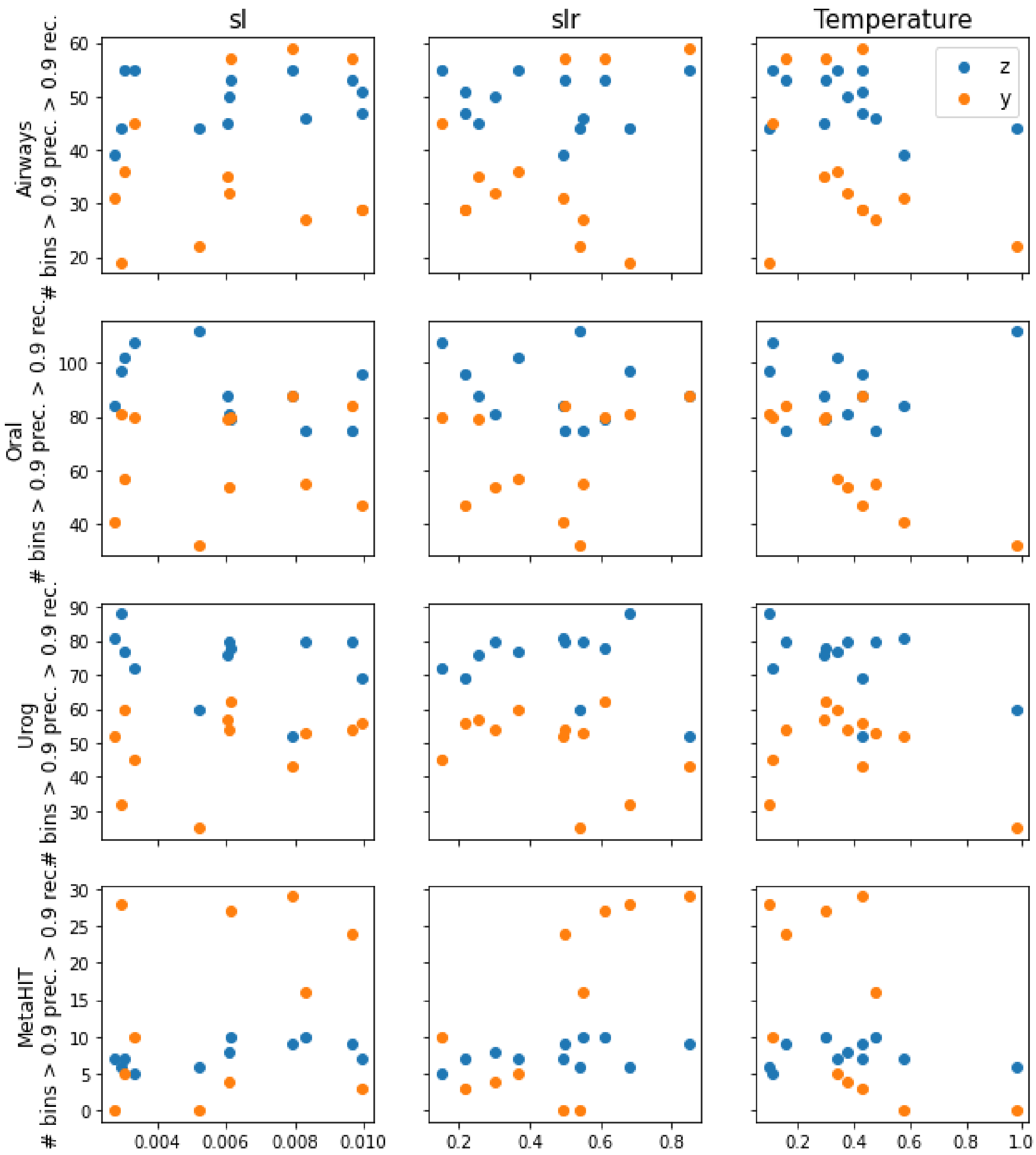


**Supplementary Figure 12. Second round of hyperparameter random searches.** AAMB(z), AAMB(y) bins above 0.9 precision and 0.9 recall are shown for Airways, Oral and Urogenital (Urog) CAMI2 datasets as well as for the MetaHIT dataset. Hyperparameters evaluated are the loss scale (sl), the discriminator loss scale (sl), and the Temperature parameter from the RelaxedOneHotCategorical distribution (Temperature). *z* , NC genomes reconstructed by the *z* latent space; *y*, NC genomes reconstructed by the *y* latent space.

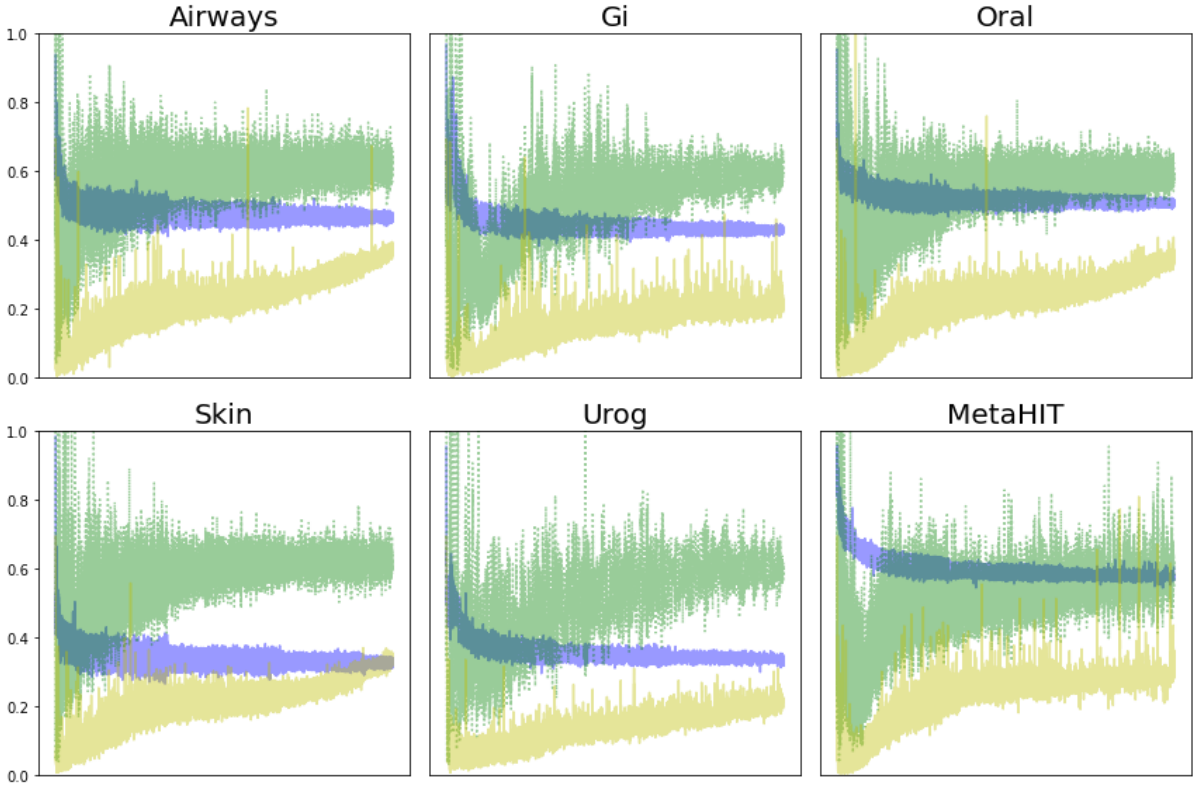


**Supplementary Figure 13. Loss along training over CAMI2 and metaHit datasets.** Adversarial autoencoder losses plots for the CAMI2 and MetaHIT datasets. Loss is plot per batch and per model. Blue: decoder, also known as generator, loss. Green: discriminator for the latent space *z* loss. Yellow: discriminator for the latent space *y* loss. Airways, CAMI2 Airways; Gi, CAMI2 Gastro intestinal; Oral, CAMI2 Oral; Skin, CAMI2 Skin; Urog, CAMI2 Urogenital.

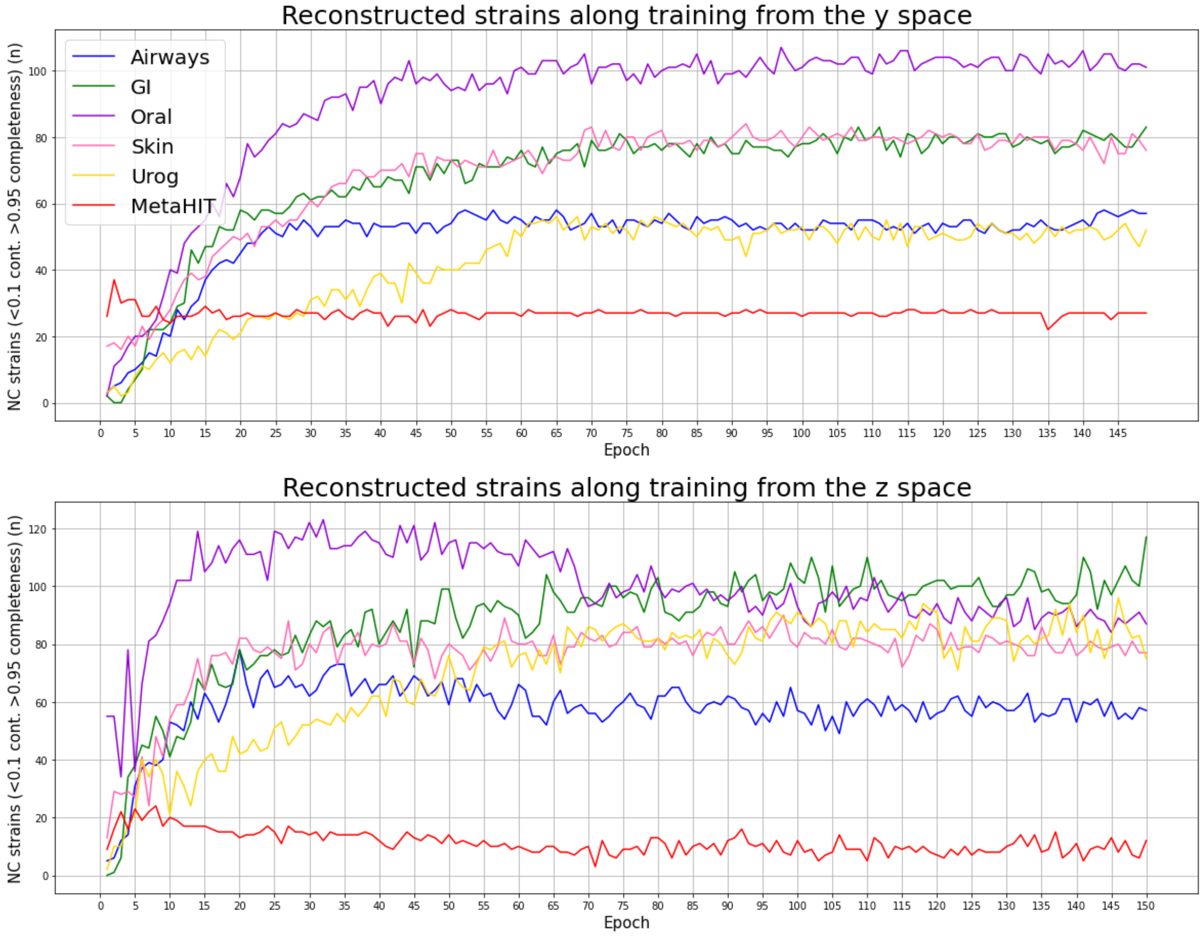


**Supplementary Figure 14. NC genomes reconstructed from the categorical (y) and continuous (z) AAMB latent spaces along 150 training epochs.** For both AAMB latent spaces, *l* and *y*, the genome reconstruction capacity was evaluated along training for the training and validation datasets. Training datasets: CAMI2 Airways, CAMI2 Oral, CAMI2 Urogenital (Urog) and MetaHIT. Validation datasets: CAMI2 Gastrointestinal (GI) and CAMI2 Skin. NC: Near complete.

| Dataset | AAMB(y) | AAMB(z) | Dereplicated | Lost |
|---|---|---|---|---|
| Airways | 34 | 69 | 74 | 7 |
| GI | 65 | 94 | 98 | 5 |
| Oral | 69 | 105 | 118 | 6 |
| Skin | 44 | 81 | 90 | 2 |
| Urog | 31 | 56 | 58 | 3 |
| MetaHIT | 29 | 11 | 22 | 7 |
| Total | 272 | 416 | 460 | 30 |

**Supplementary Table 1. dRep dereplication efficiency for AAMB near complete genomes.** Efficiency of the dereplication pipeline to dereplicate and integrate near complete genomes from AAMB(z) and AAMB(y). AAMB(y): Near complete genomes reconstructed from the y AAMB latent space. AAMB(z): Near complete genomes reconstructed from the z AAMB latent space. Dereplicated: dereplication pipeline output from the AAMB(z) and AAMB(y) NC genomes. Lost: AAMB(z) or AAMB(y) near complete genomes lost during the dereplication process. Airways: CAMI2 Airways, Oral: CAMI2 Oral, Urog: CAMI2 Urogenital, GI: CAMI2 Gastrointestinal, Skin: CAMI2 Skin.

| Dataset | AAMB(z) | AAMB(z) Y split |
|---|---|---|
| Airways | 69 | 70 |
| Gi | 94 | 97 |
| Oral | 105 | 101 |
| Skin | 81 | 82 |
| Urog | 56 | 71 |

**Supplementary Table 2. NC bins reconstructed from AAMB(z) and AAMB(z) split by their AAMB(y) label.** If our initial hypothesis that the y vector captures high-level variance and the z vector captures low-level variance, then some z clusters could be composite clusters of different y, i.e. distant organisms with similar low-level variance. However, splitting the z clusters by their y-label does not improve performance. We did hyperparameter optimisation to improve AAMB(z) y split performance, but could not get it better than AAMB(z) (data not shown). In total, this goes against our initial hypothesis.

| AAMB(z) + AAMB(y) | | | | | |
|---|---|---|---|---|---|
| Dataset | Total | dRep_CheckM2 | dRep_CheckM2 + ripping | manual dereplication | manual dereplication + ripping |
| Airways | 81 | 71 | 71 | 74 | 74 |
| Gi | 103 | 98 | 99 | 98 | 98 |
| Oral | 124 | 118 | 117 | 119 | 118 |
| Skin | 92 | 89 | 89 | 90 | 90 |
| Urog | 71 | 69 | 69 | 70 | 70 |
| MetaHIT | 29 | 15 | 15 | 22 | 22 |
| Almeida | N/A | 5078 | 5071 | 5078 | 5077 |
| HMP2 | N/A | 2723 | 2716 | 2723 | 2715 |
| AAMB(z) + AAMB(y)+VAMB | | | | | |
| Airways | 87 | 77 | 77 | 82 | 82 |
| Gi | 109 | 103 | 104 | 103 | 103 |
| Oral | 145 | 138 | 137 | 140 | 138 |
| Skin | 106 | 103 | 103 | 104 | 104 |
| Urog | 86 | 81 | 81 | 83 | 83 |
| MetaHIT | 29 | 15 | 15 | 22 | 22 |
| Almeida | N/A | 5748 | 5733 | 5748 | 5733 |
| HMP2 | N/A | 3579 | 3568 | 3573 | 3569 |

**Supplementary Table 3. Dereplication efficiency for AAMB and AVAMB near complete bins applying different dereplication strategies.** Dereplication efficiency evaluated for dRep, dRep and ripping, manual dereplication, manual dereplication and ripping. Dereplication strategies were applied to deduplicate and integrate near complete bins from AAMB y and z latent spaces, and from AAMB z, y latent spaces and VAMB. dRep: dereplication obtained by running dRep per sample. dRep + ripping: dereplication obtained by running dRep per sample and bins ripping applied subsequently. Manual dereplication: manual dereplication. Manual dereplication + ripping: manual dereplication and bins ripping applied subsequently.

| AAMB(z) + AAMB(y) | | | | |
|---|---|---|---|---|
| dataset | dRep | dRep + ripping | Manual dereplication | Manual dereplication + ripping |
| Airways | 0 | 0 | 0 | 0 |
| Gi | 3 | 0 | 3 | 0 |
| Oral | 3 | 0 | 3 | 0 |
| Skin | 0 | 0 | 0 | 0 |
| Urog | 0 | 0 | 0 | 0 |
| MetaHIT | 0 | 0 | 0 | 0 |
| Almeida | 129 | 0 | 129 | 0 |
| HMP2 | 66 | 0 | 66 | 0 |
| AAMB(z) + AAMB(y) + VAMB | | | | |
| Airways | 0 | 0 | 0 | 0 |
| Gi | 18 | 0 | 18 | 0 |
| Oral | 7 | 0 | 6 | 0 |
| Skin | 0 | 0 | 0 | 0 |
| Urog | 2 | 0 | 2 | 0 |
| MetaHIT | 0 | 0 | 0 | 0 |
| Almeida | 280 | 0 | 285 | 0 |
| HMP2 | 229 | 0 | 228 | 0 |

**Supplementary Table 4. Remaining duplicated contigs after applying different dereplication strategies to AAMB and AVAMB near complete bins.** Remaining duplicated contigs after dereplicating NC bins from AAMB and AVAMB with dRep, dRep and ripping, manual dereplication, manual dereplication and ripping. Dereplication strategies were applied to deduplicate near complete bins from AAMB z and y latent spaces and from AAMB z , y latent spaces and VAMB.. dRep: dereplication obtained by running dRep per sample. dRep + ripping: dereplication obtained by running dRep per sample and bins ripping applied subsequently. Manual dereplication: manual dereplication. Manual dereplication + ripping: manual dereplication and bins ripping applied subsequently.

| | | CPU | | | | GPU | | | | | |
|---|---|---|---|---|---|---|---|---|---|---|---|
| Dataset | BAM files parse | VAMB | | AAMB | | VAMB | | AAMB | | CheckM2 | Dereplicate |
| | | Train | Cluster | Train | Cluster | Train | Cluster | Train | Cluster | | |
| Airways | 3.3 | 27.4 | 2.1 | 35.7 | 34.5 | 6.8 | 1.2 | 10.7 | 4.8 | 210(40.6) | 2.5 |
| Gi | 4.8 | 12.2 | 0.3 | 16.5 | 0.9 | 3.35 | 0.4 | 4.7 | 0.5 | 244(48.1) | 1 |
| Oral | 4.5 | 15 | 2.5 | 38.5 | 38.3 | 7.3 | 1.2 | 11.5 | 6.4 | 250(48.5) | 2 |
| Skin | 3.8 | 26 | 1.5 | 33.4 | 23.7 | 6.6 | 1 | 10.1 | 2.4 | 242(46.3) | 2 |
| Urog | 3 | 8.9 | 0.3 | 11.2 | 0.9 | 2.8 | 0.4 | 3.4 | 0.4 | 159(32.5) | 0 |

**Supplementary Table 5. AVAMB running times both with and without GPU for all stages of the process.** The BAM files parse column indicates the contigs depths estimation runtime in minutes implemented on VAMB. AAMB, VAMB train and cluster columns represent the running times in minutes when running those models on the CAMI2 datasets with 16 CPU threads and 20 gigabytes with and without GPU. After clustering, AAMB/VAMB clusters were evaluated with Checkm2. Checkm2 was run with 8 threads and 8 Gb of memory per sample. Checkm2 runtime is expressed for all samples (average runtime per sample in minutes). Dereplication shows the runtime in minutes required to dereplicate the NC genomes generated.  All stages (bam files parsing, training, clustering, bins evaluation with checkm2 and NC bins dereplication) were integrated with snakemake for convenience and parallelization purposes. Airways: CAMI2 Airways, Oral: CAMI2 Oral, Urog: CAMI2  Urogenital, GI: CAMI2 Gastrointestinal, Skin: CAMI2 Skin.

| Dataset | AAMB(y) | AAMB(z) | VAMB | Dereplicated | Lost |
|---|---|---|---|---|---|
| Airways | 34 | 69 | 63 | 82 | 5 |
| GI | 65 | 94 | 86 | 103 | 6 |
| Oral | 69 | 105 | 124 | 138 | 6 |
| Skin | 44 | 81 | 77 | 104 | 2 |
| Urog | 31 | 56 | 78 | 83 | 4 |
| MetaHIT | 29 | 11 | 17 | 22 | 7 |
| Total | 272 | 416 | 445 | 532 | 30 |

**Supplementary Table 6. Dereplication efficiency for AAMB and VAMB near complete genomes.** Efficiency of the dereplication pipeline to dereplicate and integrate near complete genomes from AAMB(y), AAMB(z) and VAMB. AAMB(y): Near complete genomes reconstructed from the *y* AAMB latent space. AAMB(z): Near complete genomes reconstructed from the *z* AAMB latent space. VAMB: Near complete genomes reconstructed by VAMB. Dereplicated: dereplication pipeline output from the AAMB(z), AAMB(y) and VAMB NC genomes. Lost: AAMB(z), AAMB(y) or VAMB near complete genomes lost during the dereplication process. Airways: CAMI2 Airways, Oral: CAMI2 Oral, Urog: CAMI2 Urogenital, GI: CAMI2 Gastrointestinal, Skin: CAMI2 Skin.

| Dataset/model set | AAMB(z) unique | AAMB(y) unique | VAMB unique | AAMB(z) and AAMB(y) | AAMB(z) and VAMB | AAMB(y) and VAMB | AAMB(y) and AAMB(z) and VAMB | All |
|---|---|---|---|---|---|---|---|---|
| Airways | 14 | 3 | 6 | 3 | 31 | 6 | 19 | 82 |
| Gi | 13 | 3 | 5 | 4 | 22 | 5 | 51 | 103 |
| Oral | 12 | 5 | 18 | 2 | 41 | 14 | 47 | 139 |
| Skin | 22 | 2 | 14 | 3 | 25 | 8 | 30 | 104 |
| Urog | 5 | 3 | 13 | 0 | 33 | 1 | 28 | 83 |
| MetaHIT | 0 | 6 | 0 | 1 | 0 | 6 | 9 | 22 |

**Supplementary Table 7. VAMB AAMB NC genomes contributions and intersections.** AAMB and VAMB specific contributions and intersections on CAMI2 and MetaHIT datasets. Unique stands for NC genomes only reconstructed by the given model. And stands for NC genomes reconstructed by models and not any of the rest. All presents the addition of all sets which at the same time expresses the total amount of NC genomes reconstructed by VAMB and AAMB when dereplicating with dRep. Airways: CAMI2 Airways, Oral: CAMI2 Oral, Urog: CAMI2 Urogenital, GI: CAMI2 Gastrointestinal, Skin: CAMI2 Skin. NC: Near complete.

| Dataset | A | V | M | V+A | V+M | A+M | V+M+A |
|---|---|---|---|---|---|---|---|
| Airways | **74** | 63 | 36 | **82** | 61 | 77 | 84 |
| GI | **98** | 86 | 76 | 103 | 101 | **108** | 112 |
| Oral | 118 | **124** | 68 | **138** | 129 | 128 | 145 |
| Skin | **90** | 72 | 62 | **104** | 97 | 98 | 111 |
| Urog | 70 | **78** | 66 | 83 | **88** | **88** | 96 |
| MetaHIT | **22** | 17 | 1 | **22** | 13 | 21 | 22 |
| Total | **472** | 440 | 309 | **532** | 489 | 520 | 570 |

**Supplementary Table 8. NC genomes reconstructed from the benchmark datasets by AAMB, VAMB, MetaBAT2 individually and when integrating with the dereplication pipeline.** Near complete genomes reconstructed by AAMB, VAMB, MetaBAT2, VAMB and AAMB ,VAMB and MetaBAT2**,** AAMB and MetaBAT2, VAMB and AAMB and MetaBAT2. Integration of binning results was done with the dereplication pipeline. A: AAMB, V: VAMB, M: MetaBAT2, V+A: VAMB and AAMB, V+M: VAMB and MetaBAT2, A+M: AAMB +MetaBAT2,  V+A+M: VAMB and AAMB and MetaBAT2.

| Dataset | AAMB | VAMB | MetaBAT2 | AVAMB | SemiBin |
|---|---|---|---|---|---|
| Airways | 74 | 63 | 36 | 82 | 96 |
| GI | 98 | 86 | 76 | 103 | 140 |
| Oral | 118 | 124 | 68 | 138 | 159 |
| Skin | 90 | 72 | 62 | 104 | 138 |
| Urog | 70 | 78 | 66 | 83 | 112 |
| Total | 472 | 440 | 309 | 532 | 645 |

**Supplementary Table 9. NC genomes reconstructed from the CAMI2 datasets by AAMB, VAMB, MetaBAT2, AVAMB and SemiBin.** Near complete genomes reconstructed by AAMB, VAMB, MetaBAT2, AVAMB and SemiBin.

| Dataset | Airways | GI | Oral | Skin | Urog |
|---|---|---|---|---|---|
| Runtime | 18.7 | 13.2 | 20.01 | 14.05 | 9.6 |

**Supplementary Table 10. SemiBin running time on GPU for the CAMI datasets.** SemiBin running times in hours when running on 20 CPU threads and 150 gigabytes with one GPU on the CAMI benchmark datasets. GI ; Gastrointestinal, Urog; Urogenital.

| Dataset | V = A | V > A | A > V | V unique | A unique | AVAMB |
|---|---|---|---|---|---|---|
| Almeida | 332 | 1530 | 2091 | 670 | 1110 | 5733 |
| HMP2 | 62 | 961 | 782 | 848 | 916 | 3569 |

**Supplementary Table 11. VAMB and AAMB bins from the same sample with 100% identity over at least 75% of the smallest bin**. NC: Near complete. V = A: VAMB and AAMB NC bins with exact same score. V > A: VAMB NC bins with higher score than AAMB NC bins, A > V: AAMB NC bins with higher score than VAMB NC bins, V uniq: VAMB NC bins not reconstructed by any AAMB NC bin at the selected identity settings, A unique: AAMB NC bins not reconstructed by any VAMB NC bin at the selected identity settings.

| Level | Common | VAMB only | AAMB only | AVAMB |
|---|---|---|---|---|
| Domain | 2 | 0 | 0 | 2 |
| Phylum | 13 | 0 | 0 | 13 |
| Class | 16 | 0 | 0 | 16 |
| Order | 38 | 2 | 1 | 41 |
| Family | 81 | 5 | 3 | 89 |
| Genus | 286 | 22 | 18 | 325 |
| Species | 614 | 63 | 93 | 767 |

**Supplementary Table 12. We annotated all Almeida NC bins dereplicated bins from AAMB and VAMB with GTDB-Tk and counted a particular taxon if at least one genome was reconstructed.** Common column presents the amount of taxons with at least one NC bin generated both by VAMB and AAMB, the VAMB only column presents the amount of taxons with at least one NC bin generated only by VAMB, and the AAMB only column presents the amount of taxons with at least one NC bin generated only by AAMB. Worth to mention that the quantities do not reflect how many NC bins were reconstructed for each taxon. Results also represented in Figure 2c.

| Level | Common | VAMB only | AAMB only | AVAMB |
|---|---|---|---|---|
| Domain | 2 | 0 | 0 | 2 |
| Phylum | 12 | 0 | 0 | 12 |
| Class | 14 | 0 | 0 | 14 |
| Order | 27 | 1 | 0 | 28 |
| Family | 47 | 3 | 1 | 51 |
| Genus | 122 | 8 | 18 | 148 |
| Species | 200 | 17 | 61 | 278 |

**Supplementary Table 13. We annotated all Human Microbiome Project 2 NC dereplicated bins from AAMB and VAMB with GTDB-Tk and counted a particular taxon if at least one genome was reconstructed.** Common column presents the amount of taxons with at least one NC bin generated both by VAMB and AAMB, the VAMB only column presents the amount of taxons with at least one NC bin generated only by VAMB, and the AAMB only column presents the amount of taxons with at least one NC bin generated only by AAMB. Worth to mention that the quantities do not reflect how many NC bins were reconstructed for each taxon. Results also represented in Supplementary Figure 14.

| Dataset | Parse bam files | VAMB | | AAMB | | CheckM2 | dereplicate |
|---|---|---|---|---|---|---|---|
| | | Train | Cluster | Train | Cluster | | |
| Almeida | 1.4 | 5.32 | 1.6 | 7.22 | 6.8 | 0.25(0.07) | 0.6 |

**Supplementary Table 14. AVAMB running times when running with GPU on a high-performance computing station for the Almeida dataset.** The BAM files parse column indicates the contigs depths estimation runtime in hours implemented on VAMB. AAMB, VAMB train and cluster columns represent the running times in hours when running those models on the Almeida datasets with 20 CPU threads, 1 GPU, and 100 GB RAM. After clustering, AAMB/VAMB clusters were evaluated with Checkm2. Checkm2 was run on parallel per sample, assigning 25 threads and 40 Gb of memory per sample run. Checkm2 runtime expressed the time spent for all samples (average runtime per sample in hours). Dereplication shows the runtime in hours required to dereplicate the NC genomes generated. All stages (bam files parsing, training, clustering, bins evaluation with checkm2 and NC bins dereplication) were integrated with snakemake for convenience and parallelization purposes.

| Dataset | A | V | M | V+A | V+M | A+M | V+M+A |
|---|---|---|---|---|---|---|---|
| Almeida | 5077 | 4617 | 2954 | **5733** | 5419 | 5721 | 6121 |

**Supplementary Table 15. NC bins reconstructed from the Almeida dataset by AAMB, VAMB, MetaBAT2 individually and when integrating with dRep.** Near complete bins reconstructed by AAMB, VAMB, MetaBAT2, VAMB and AAMB ,VAMB and MetaBAT2**,** AAMB and MetaBAT2, VAMB and AAMB and MetaBAT2. Integration of binning results was done with dRep. A: AAMB, V: VAMB, M: MetaBAT2, V+A: VAMB and AAMB, V+M: VAMB and MetaBAT2, A+M: AAMB +MetaBAT2, V+A+M: VAMB and AAMB and MetaBAT2.

| Dataset | AAMB(z) | AAMB(y) | VAMB | AAMB(z+y) | AAMB+VAMB |
|---|---|---|---|---|---|
| Almeida | 4,355 | 2,891 | 4,630 | 5,077 | 5,733 |
| HMP2 | 2,586 | 623 | 2,655 | 2,715 | 3,569 |

**Supplementary Table 16. NC bins reconstructed from the Almeida and Human Microbiome Project 2 datasets by AAMB(z), AAMB(y), VAMB, individually and when integrated.** Near complete genomes reconstructed by AAMB(z), AAMB(y), VAMB, AAMB(z) and AAMB(y), VAMB and AAMB(z) and AAMB(y). Integration of binning results was done with the dereplication pipeline. AAMB(z): NC bins reconstructed from AAMB z latent space, AAMB(y): NC bins reconstructed from AAMB y latent space, AAMB(z+y): NC bins reconstructed from AAMB z latent space and AAMB y latent space, AAMB + VAMB: NC bins reconstructed from AAMB z latent space and AAMB y latent space and VAMB.

| Dataset | V = A | V > A | A > V | V unique | A unique | AVAMB |
|---|---|---|---|---|---|---|
| Airways | 41 | 22 | 43 | 20 | 47 | 162 |
| GI | 101 | 24 | 35 | 12 | 41 | 199 |
| Oral | 83 | 67 | 69 | 45 | 46 | 293 |
| Skin | 55 | 21 | 38 | 19 | 40 | 168 |
| Urog | 59 | 14 | 24 | 30 | 21 | 142 |
| MetaHIT | 0 | 3 | 13 | 0 | 12 | 22 |

**Supplementary Table 17. VAMB and AAMB bins from the same sample with 100% identity over at least 75% of the smallest bin scores comparison**. NC: Near complete. V = A: VAMB and AAMB NC bins with exact same score. V > A:  VAMB NC bins with higher score than AAMB NC bins, A > V: AAMB NC bins with higher score than VAMB NC bins, V uniq: VAMB NC bins not reconstructed by any AAMB NC bin at the selected identity settings, A unique: AAMB NC bins not reconstructed by any VAMB NC bin at the selected identity settings.
